# Sphingosine-1-phosphate signaling through Müller glia regulates neuroprotection and the accumulation of immune cells in the rodent retina

**DOI:** 10.1101/2025.02.03.636254

**Authors:** Olivia Taylor, Lisa Kelly, Heithem El-Hodiri, Andy J. Fischer

**Author notes:** **Corresponding author**: Andy J. Fischer, Department of Neuroscience, Ohio State University, College of Medicine, 3020 Graves Hall, 333 W. 10^th^ Ave, Columbus, OH 43210-1239, USA.

## Abstract

The purpose of this study was to investigate how Sphingosine-1-phosphate (S1P) signaling regulates glial phenotype, neuroprotection, and reprogramming of Müller glia (MG) into neurogenic MG-derived progenitor cells (MGPCs) in the adult mouse retina. We found that S1P-related genes were dynamically regulated following retinal damage. *S1pr1* (encoding S1P receptor 1) and *Sphk1* (encoding sphingosine kinase 1) are expressed at low levels by resting MG and are rapidly upregulated following acute damage. Overexpression of the neurogenic bHLH transcription factor Ascl1 in MG downregulates *S1pr1*, and inhibition of Sphk1 and S1pr1/3 enhances Ascl1-driven differentiation of bipolar-like cells and suppresses glial differentiation. Treatments that activate S1pr1 or increase retinal levels of S1P initiate pro-inflammatory NFκB-signaling in MG, whereas treatments that inhibit S1pr1 or decreased levels of S1P suppress NFκB-signaling in MG in damaged retinas. Conditional knock-out of NFκB-signaling in MG increases glial expression of *S1pr1* but decreases levels of *S1pr3* and *Sphk1*. Conditional knock-out (cKO) of *S1pr1* in MG, but not *Sphk1*, enhances the accumulation of immune cells in acutely damaged retinas. cKO of *S1pr1 i*s neuroprotective to ganglion cells, whereas cKO of *Sphk1* is neuroprotective to amacrine cells in NMDA-damaged retinas. Consistent with these findings, pharmacological treatments that inhibit S1P receptors or inhibit Sphk1 had protective effects upon inner retinal neurons. We conclude that the S1P-signaling pathway is activated in MG after damage and this pathway acts secondarily to restrict the accumulation of immune cells, impairs neuron survival and suppresses the reprogramming of MG into neurogenic progenitors in the adult mouse retina.

## Introduction

Neuronal damage and inflammation impact the ability of Müller glia (MG) to become proliferating, neurogenic progenitor-like cells in different vertebrate models of retinal regeneration. The Nuclear Factor Kappa B (NFκB) cell signaling pathway is a key mediator of inflammation that induces the reactivity of MG, acts to restore glial quiescence and suppress the neurogenic potential of MG-derived progenitors in the mouse retina (Hoang et al., 2020; Palazzo et al., 2022). In acutely damaged retinas, NFκB-signaling is selectively activated in MG by pro-inflammatory cytokines that are produced by activated microglia (Palazzo et al., 2022; Palazzo et al., 2023). Inhibition of NFκB in MG results in diminished recruitment of immune cells into damaged retinas, increased neuronal survival, and increased formation of neuron-like cells with forced expression of the pro-neural basic Helix-Loop-Helix (bHLH) transcription factor Ascl1 (Palazzo et al., 2022). NFκB is known to be coordinated with Sphingosine-1-phosphate (S1P) -signaling to regulate different cellular processes including inflammation (Blom et al., 2010; Pérez-Jeldres et al., 2021; Zheng et al., 2019).

S1P is aliphatic amino alcohol that is synthesized by sphingosine kinases (SPHK1 and SPHK2) and degraded by a lyase (SGPL1) (Fig. 1). S1P is exported from cells by transporters (MFSD2A and SPNS2) or hydrolyzed back to sphingosine by a phosphatase (SGPP1) (Fig. 1). Ceramide and S1P metabolism are highly coordinated: sphingosine is produced by ASAH1 via hydrolysis of ceramide or modified by ceramide synthases (CERS2/5/6) that catalyze the transfer of an acyl chain from acyl-CoA (Fig. 1). Secreted S1P is chaperoned by HDL-anchored ApoM or albumin and binds to and activates G-protein coupled receptors (S1PR1-S1PR5) to elicit a wide array of functions across different cell types (Fig. 1) (Murata et al., 2000; Obinata and Hla, 2019). S1PRs activate different second messenger pathways including MAPK, PI3K/mTor and Jak/Stat, and crosstalk with pro-inflammatory pathways such as NFκB (Fig. 1) (Gurgui et al., 2010; Hu et al., 2020). S1P-signaling is known to mediate inflammatory responses, cellular proliferation, cell survival and angiogenesis in lymphocytes and endothelial cells (Obinata and Hla, 2019). In the developing retina, S1P-signaling is required for vascular maturation, progenitor cell cycle exit, and axon guidance (Simón et al., 2019). S1P can also have negative effects on the retina which include migration of MG, neovascularization, and inflammation associated with proliferative retinopathies and aging (Shiwani et al., 2021). Many of these studies applied clinically relevant S1P analogs, which target S1P receptors and ceramide pathway components nonspecifically, thereby complicating the interpretation of findings. Additionally, most of these studies on retinal cells were conducted *in vitro* and need to be followed-up by *in vivo* analyses to identify the coordinated impact of S1P-signaling on the many different types of retinal cells.

**Figure 1:**
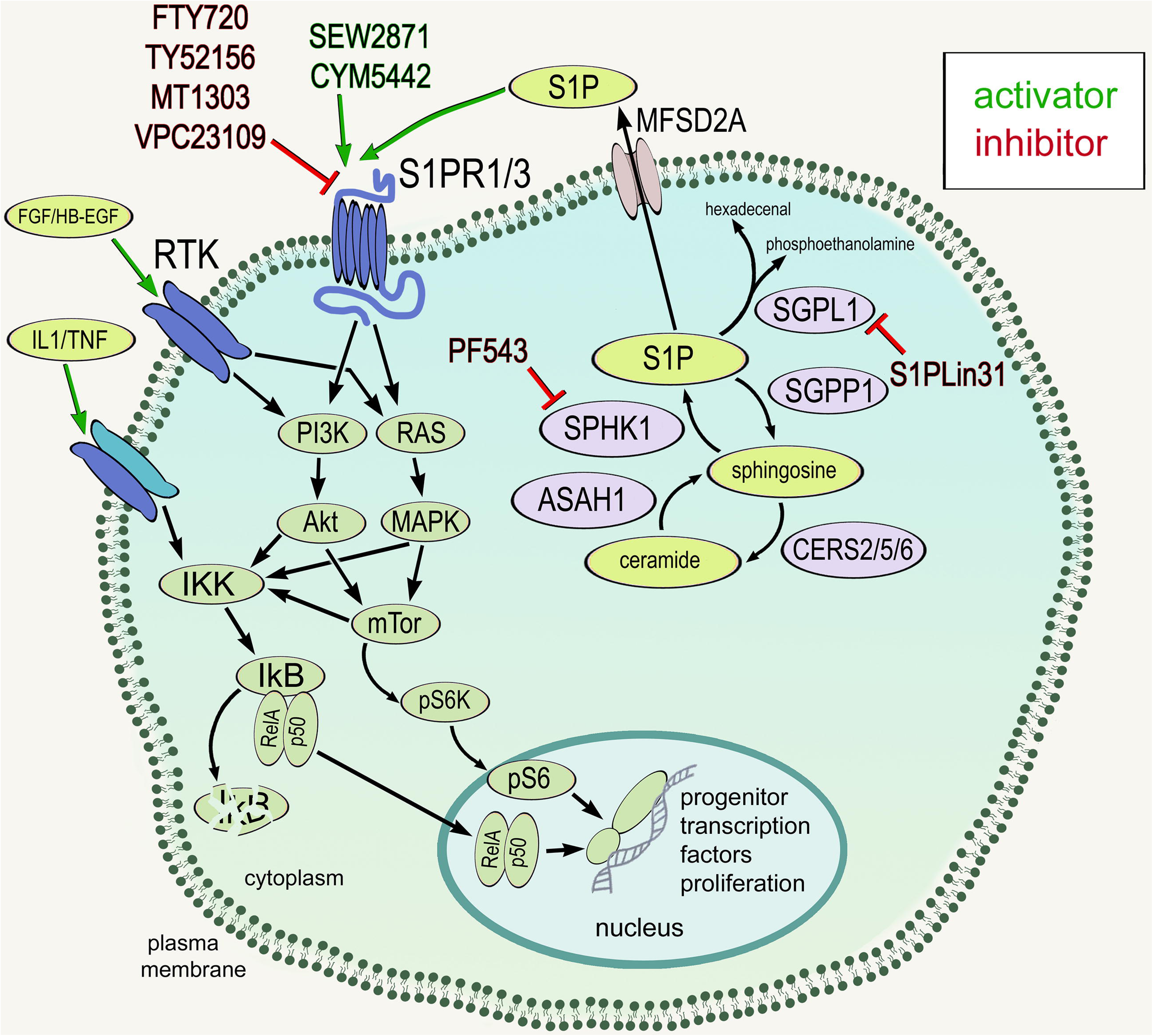
Schematic summary of S1P synthesis, degradation, cell signaling and sites of action of different agonists and antagonists. Red lines indicate inhibitors or antagonists and green arrows indicate activators or agonists. Abbreviations: S1P, sphingosine 1-phosphate; S1pr, sphingosine 1 phosphate receptor; Sphk1, sphingosine kinase-1; ASAH1, acid ceramidase 1; CERS, ceramide synthase; SGPP1, sphingosine 1-phosphate phosphatase 1.

We recently identified the effects of S1P-signaling in the avian retina, wherein treatments that inhibited S1pr1 or decreased levels of S1P enhanced MGPC formation and neurogenesis in damaged retinas (Taylor et al., 2024b). S1P-related genes are highly expressed by neurons and glia, and levels of expression are dynamically regulated following damage to the chick retina. *S1PR1* was highly expressed by resting MG and is rapidly downregulated following damage. This pattern of expression is similar to that seen in zebrafish and human retinas. Further, ablation of microglia from damaged retinas, wherein the formation of MGPCs is blocked (Fischer et al., 2014), has a significant impact upon expression patterns of S1P-related genes in MG, and this may, in part, be mediated by TGFβ/Smad3-signaling which maintains high levels S1PR1 expression in resting MG (Taylor et al., 2024b). Thus, in the chick retina, S1P-signaling is dynamically regulated in MG to suppress MGPC formation and activation of S1P-signaling depends, in part, on signals produced by activated microglia.

The neuroprotective effects of S1P receptor-targeting drugs have been reported previously (Basavarajappa et al., 2023b; Nakamura et al., 2021), but the mechanisms underlying neuroprotection is not well understood. Further, S1P-signaling has not been targeted with a tissue-specific manner to assess neuroprotection and inflammation. Accordingly, the purpose of this study was to better understand how S1P-signaling impacts inflammation, glial reactivity, neuronal survival and reprogramming of MG into neurogenic MGPCs in the mouse retina.

## Methods

### Animals

The animal use approved in these experiments was in accordance with the guidelines established by the National Institutes of Health and IACUC at The Ohio State University. Mice were housed under a 12/12 light-dark cycle and received water and chow ad libitum. NFκB-eGFP reporter mice, which have eGFP-expression driven by a chimeric promoter containing three HIV-NFκBcis elements (Magness et al., 2004). and *Ikkb^fl/fl^* mice, with insertion of Cre recombinase binding sites (LoxP) into the intronic regions flanking exon 3 of the wild type *Ikkb* gene (Li et al., 2003). were kindly provided Dr. Denis Guttridge’s laboratory at The Ohio State University. We crossed *Sphk1^fl/fl^* and *S1pr1^fl/fl^* mice (provided by Dr. Timothy Hla; Harvard Medical School) onto *Rlbp1-CreERT;R26-stop-flox-CAG-tdTomato* mice (provided by Dr. Ed Levine; Vanderbilt University), wherein Cre-mediated recombination occurs in a tamoxifen-dependent manner specifically in MG under the control of the retinaldehyde binding protein 1 (*Rlbp1*) promoter, herein referred to as *Rlbp1-CreERT:S1pr1^fl/fl^* and *Rlbp1-CreERT:Sphk1^fl/fl^*. The use of Ascl1 over-expressing mice (*Glast-CreER:LNL-tTA:tetO-mAscl1-ires-GFP)* was as previously described (Palazzo et al., 2022; Ueki et al., 2015).

### Ablation of microglia

Mice were provided control or PLX5622 (1200 ppm) (Research Diets #D1100404i) diet *ad libitum* for two weeks to deplete microglia prior to experimental paradigms. We and other groups have validated the efficacy of this treatment paradigm to deplete microglia in the mouse retina (Palazzo et al., 2022; Todd et al., 2020).

### Intraocular Injections

Mice were anesthetized via inhalation of 2.5% isoflurane in oxygen and intraocular injections performed as described previously (Hamilton #7803-05 10 MM 12DEG) (Palazzo et al., 2022). For all experiments, the vitreous chamber of right eyes of mice were injected with the experimental compound and the contra-lateral left eyes were injected with a control vehicle. Injections of 100 mM NMDA in sterile saline were administered at a volume of 1 ul. Compounds included in this study are described in Table 1. For adult mice, the average eye weight is 0.02 grams. Doses were estimated from common mg/kg doses reported for oral administration. Injection paradigms are included in each figure. Intraperitoneal injections of tamoxifen (Sigma T5648; 1.5 mg/100 l corn oil per dose) were performed for 4 consecutive days to induce ER-Cre activity.

**Table 1.**
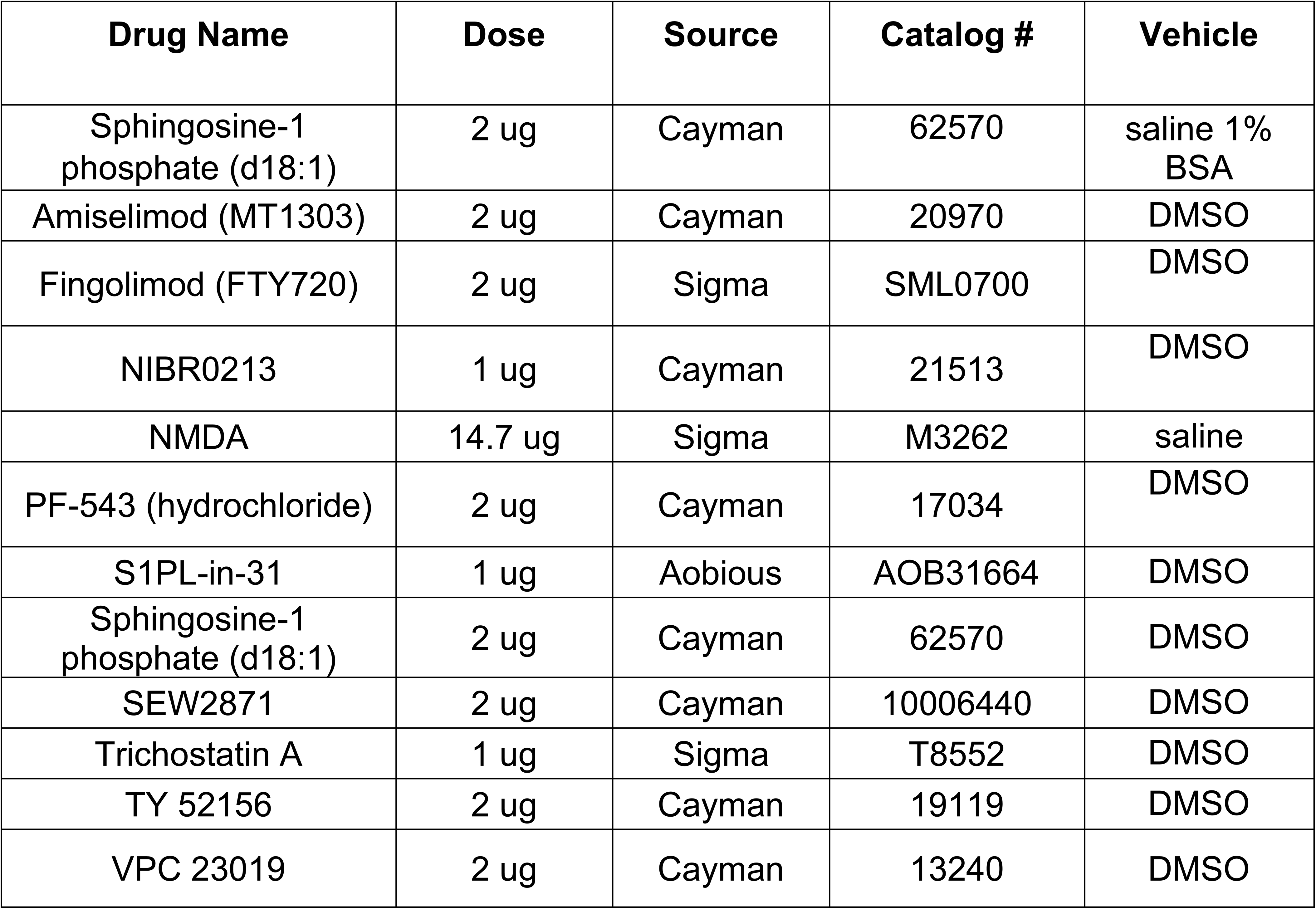
Pharmacological Compounds.

### Fixation, sectioning, and immunocytochemistry

Retinas were fixed, sectioned, and immunolabeled as described previously (Fischer et al., 1998). To identify MG, we labeled sections for Sox2, which is known to label the nuclei or MG in the INL and the nuclei of astrocytes at the vitread surface of the retina (Fischer et al., 2010). To identify CNS microglia/macrophages and peripheral monocyte-derived macrophages, we labeled sections for Iba1 and CD45, as described previously (Palazzo et al., 2022). None of the observed labeling was due to non-specific labeling of secondary antibodies or auto-fluorescence because sections labeled with secondary antibodies alone were devoid of fluorescence. Primary antibodies used in this study are described in **Table 2**. Secondary antibodies included donkey-anti-goat-Alexa488/594/647 (Life Technologies A11055; A11058; A21447), donkey-anti-rabbit-Alexa488/594 (Life Technologies A21206; A21207); and goat-anti-mouse-Alexa488/568 (Life Technologies A11001; A-11004) diluted to 1:1000 in PBS plus 0.2% Triton X-100. Nuclear staining was accomplished using DRAQ5 (Thermo 62251) or DAPI (Sigma D9542).

**Table 2:** Antibody table.

| **Antibody** | **Dilution** | **Host** | **Source** | **Clone/Catalog number** |
| --- | --- | --- | --- | --- |
| Brn3a | 1:50 | Mouse | Millipore | MAB1585 |
| CD45 | 1:300 | Rat | BioRad | MCA1031G |
| cFos | 1:500 | Rabbit | Cell Signaling | 2250 |
| GFP | 1:1000 | Chicken | Abcam | Ab13970 |
| HuD/C | 1:300 | Mouse | Invitrogen | A21271 |
| Iba1 | 1:500 | Rabbit | Wako Pure Chemical Industries | 019–19741 |
| PAX6 | 1:1000 | Rabbit | Biolegend | 901301 |
| pERK1/2 | 1:500 | Rabbit | Cell Signaling | 4370 |
| pS6 | 1:500 | Rabbit | Cell Signaling | 2215 |
| Sox2 | 1:1000 | Goat | R&D Systems | KOY0418121 |
| S1pr1 |  | Rabbit | Novus Biologicals | NB120-11424 |
| S1pr1 |  | Mouse | R&D Systems | MAB7089 |
| S1pr1 |  | Mouse | Santa Cruz | Sc-48356 |
| Sphk1 |  | Rabbit | Bioss Antibodies | bs-2652R |
| Sphk1 |  | Rabbit | Cell Signaling | 12071 |

### Labeling for EdU

Regular drinking water was removed 24hr prior to NMDA injections and replaced with sterile water containing 5-ethynyl-2′-deoxyuridine (EdU; Sigma; 50 mg/100 mL dH_2_O) which was replaced every third day. Mice were maintained on EdU water until the fourth day after TSA treatment. For the detection of nuclei that incorporated EdU, immunolabeled sections were fixed in 4% formaldehyde in 0.1M PBS pH 7.4 for 5 minutes at room temperature. Samples were washed for 5 minutes with PBS, permeabilized with 0.5% Triton X-100 in PBS for 1 minute at room temperature and washed twice for 5 minutes in PBS. Sections were incubated for 30 minutes at room temperature in a buffer consisting of 100 mM Tris, 8 mM CuSO_4_, and 100 mM ascorbic acid in dH_2_O. Alexa Fluor 647 Azide (Thermo Fisher Scientific #A10277) was added to the buffer at a 1:1000 dilution.

### Terminal deoxynucleotidyl transferase dUTP nick end labeling (TUNEL)

The TUNEL method was used to identify dying cells with fragmented DNA. An *in-situ* Cell Death Detection kit from Roche (Fluorescein, #11684795910) was used according to manufacturer instructions.

### Fluorescent in situ hybridization

Standard procedures were used for fluorescent *in-situ* hybridization (FISH), as described previously (Campbell et al., 2021a; Taylor et al., 2024a). In short, retinas from P9 eyes were fixed for 4 hours RT in 4% paraformaldehyde buffered in 0.1M dibasic sodium phosphate, washed in PTW (PBS + 0.2% Tween), and incubated in 30% sucrose at 4°C overnight. The retinas were embedded in OCT-compound and cryo-sectioned at 12 microns. Tissue sections were processed for in situ hybridization with a split-initiator probe pair (Molecular Instruments) according to the manufacturer protocol for fresh/fixed frozen tissues. For slides in which immunocytochemistry was conducted with FISH, primary antibodies incubated overnight with the hairpin amplification buffer solution, and secondary antibodies incubated for one hour the next day. Slides were mounted with glycerol and glass coverslips.

### Photography, immunofluorescence measurements, and statistics

Wide-field photomicroscopy was performed using a Leica DM5000B microscope equipped with epifluorescence and Leica DC500 digital camera. Confocal images were obtained using a Leica SP8 imaging system at the Department of Neuroscience Imaging Facility at the Ohio State University. Images were optimized for color, brightness and contrast, multiple channels over laid and figures constructed using Adobe Photoshop. Cell counts were performed on representative images. To avoid the possibility of region-specific differences within the retina, cell counts were consistently made from the same region of retina for each data set.

Counts of FISH puncta per cell were made as follows. The number of FISH puncta per cell was determined for fixed regions of interest (ROI) of the INL by counting the total number of FISH puncta and dividing by the total number of Sox2^+^ MG per ROI. The number of FISH puncta per transgenic cell was determined by counting the number of FISH puncta withing GFP^+^ cells divided by the number of Sox2^+^GFP^+^ MG within the ROI. The number of FISH puncta per wild type (WT) cell was determined by counting the number of FISH puncta outside of GFP^+^ cells and dividing by the number of Sox2^+^GFP^-^ MG within the ROI. Image J was used to establish threshold fluorescence intensity levels to capture GFP^+^ cells and FISH puncta, and for water-shedding and counting numbers of FISH puncta.

Where significance of difference was determined between two treatment groups accounting for inter-individual variability (means of treated-control values) we performed a two-tailed, paired t-test. Where significance of difference was determined between two treatment groups, we performed a two-tailed, unpaired t-test. Significance of difference between multiple groups was determined using one way ANOVA followed by Sidek correction. GraphPad Prism 10 was used for statistical analyses and generation of histograms and bar graphs.

### scRNA-seq

We analyzed scRNA-seq libraries that were generated and characterized previously (Hoang et al., 2020; Li et al., 2024; Palazzo et al., 2022). For WT retinas, dissociated retinal cells were loaded onto the 10X Chromium Cell Controller with Chromium 3’ V2 reagents. Using Seurat toolkits (Powers and Satija, 2015; Satija et al., 2015), Uniform Manifold Approximation and Projection (UMAP) for dimensional reduction plots were generated from 9 separate cDNA libraries, including 2 replicates of control undamaged retinas, and retinas at different times after NMDA-treatment (Hoang et al., 2020). In the IKK cKO whole retina library, dissociated cells were loaded onto the 10X Chromium Cell Controller with Chromium 3’ V3 reagents. Using Seurat toolkits, UMAP for dimensional reduction plots were generated from aggregated cDNA libraries, including 2 replicates of undamaged×WT, damaged×WT, undamaged×IKK cKO, and damaged×IKK cKO retinas (Palazzo et al., 2022). In both groups of libraries, Seurat was used to construct gene lists for differentially expressed genes (DEGs), violin/scatter plots, and dot plots. Significance of difference in violin/scatter plots was determined using a Wilcoxon Rank Sum or Poisson tests with Bonferroni correction. Genes that were used to identify different types of retinal cells included the following: (1) Müller glia: *Glul, Vim, Scl1a3, Rlbp1*, (2) MGPCs: *PCNA, CDK1, TOP2A, ASCL1*, (3) microglia: *C1qa, C1qb, Ccl4, Csf1r*, (4) ganglion cells: *Thy1, Pou4f2, Rbpms2, Nefl*, (5) amacrine cells: *Gad67, Calb2, Tfap2a*, (6) horizontal cells: *Prox1, Calb2, Ntrk1*, (7) bipolar cells: *Vsx1, Otx2, Grik1, Gabra1*, (7) cone photoreceptors: *Gnat2, Gnb3, Opn1lw*, (8) rod photoreceptors: *Rho, Nr2e3, Arr3,* (9) astrocytes: *Pax2, S100b*, *Gja1*, and (10) endothelial cells: *Pecam1, Cdh5, Tie1*.

## Results

### Expression of S1P-related genes in the retina

We began by analyzing patterns of expression of S1P-related genes in a large integrated scRNA-seq database of mouse retinal cells from different postnatal ages ranging from 2 to 25 weeks; this database has been generated and described previously (Li et al., 2024). This aggregated database contained more than 330,000 cells which form numerous distinct clusters of retinal cells in Uniform Manifold Approximation and Projection (UMAP) plots (Figs. 2a). Glial cells formed distinct clusters based on patterns of expression of cell-distinguishing markers (see Methods). For example, we found *Rlbp1* in MG (and RPE), *Pax2* in astrocytes and *C1qa* in microglia (Fig. 2b). We next examined patterns of expression and changes in expression levels of S1P-related genes (Figs. 2c-f). *Sphk1* was detected in astrocytes, MG and a few clusters of amacrine cells (Fig. 2c). By comparison, *S1pr1* and *S1pr3* were detected at high levels in MG (Figs. 2d,e). Additionally, *S1pr1* was detected in endothelial cells and *S1pr3* was detected at high levels in pericytes and at lower levels in endothelial cells and astrocytes (Figs. 2d,e). *Sgpl1* was expressed at high levels in most microglia and in cells scattered across all different major cell types (Fig. 2f).

**Figure 2:**
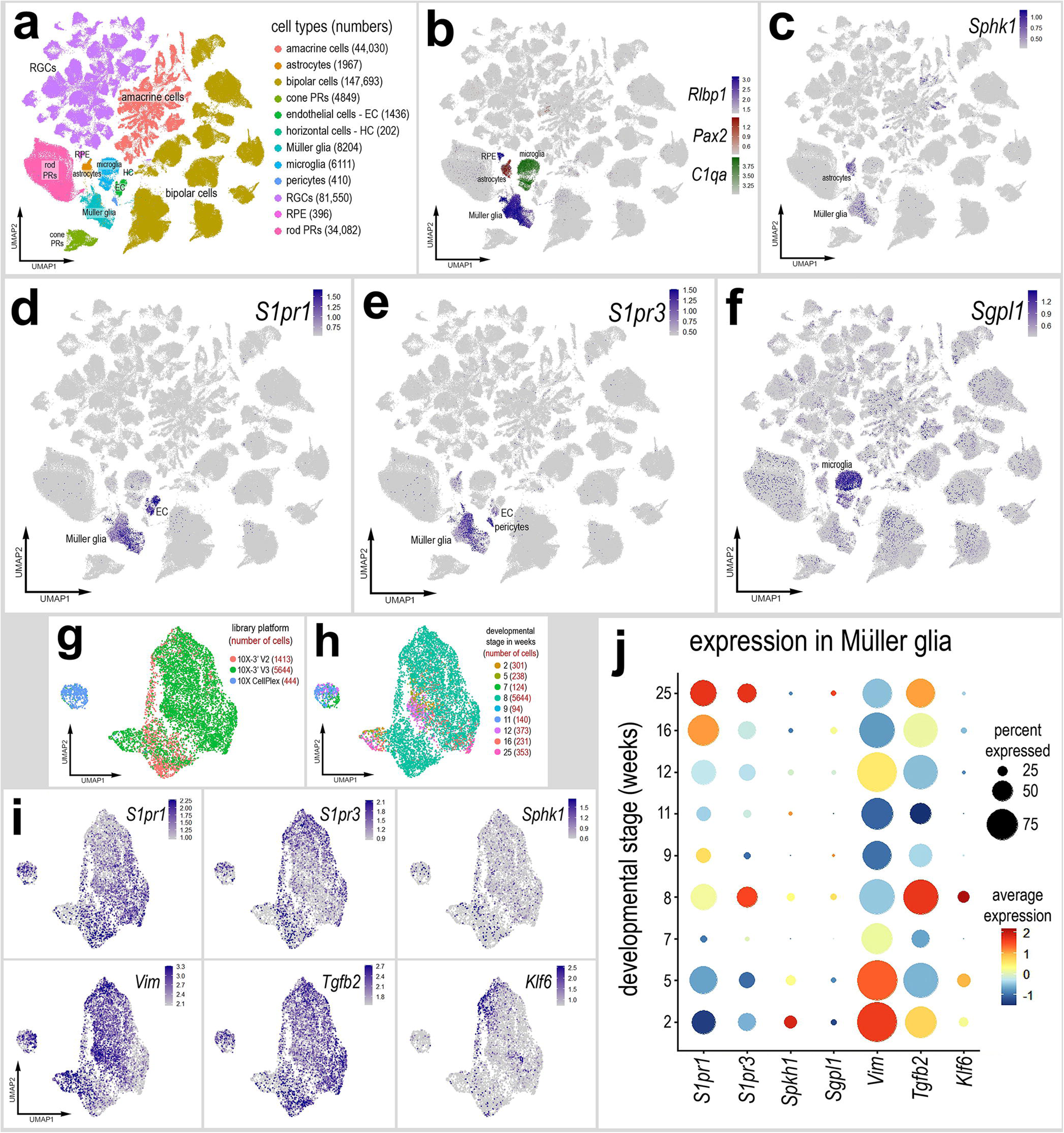
Patterns of expression of S1P-related genes in the mouse retina at different postnatal ages. scRNA-seq was used to identify patterns of expression of S1P-related factors among retinal cells with the data presented in UMAP (**a-i**) or dot plot (**j**). Aggregate scRNA-seq libraries were generated for retinal cells at 9 stages of postnatal development ranging from 2 to 25 weeks of age. UMAP-ordered cells formed distinct clusters of RPE, MG, astrocytes, and microglia based on distinct patterns of gene expression (**b**). UMAP heatmap plots illustrate patterns and levels of expression of *Sphk1* (**c**), *S1pr1* (**d**), *S1pr3* (**e**) and *Sgpl1* (**f**). MG were bioinformatically isolated and analyzed from the aggregate library (**g-j**). UMAP heatmap plots and dot plot illustrate levels of expression of S1P-related genes (*S1pr1*, *S1pr3*, and *Sphk1*) and reactivity-associated genes (*Vim*, *Tgfb2*, and *Klf6)* (**i**, **j**). Abbreviations: MG, Müller glia; NMDA, N-methyl-D-aspartate; RPE, retinal pigmented epithelium; UMAP, uniform manifold approximation and projection.

We next bioinformatically isolated the MG to perform more detailed analyses of S1P-related genes. We re-normalized and re-established principal components for UMAP embedding, filtered clusters with abnormally high features/UMI per cell, filtered contamination from astrocytes (high *Pax2* and *S100b*), and bipolar cells (high *Lhx4, Otx2, Scgn*). These processes produced distinct clusters of MG, however these cells were segregated by platform, but not developmental stage (Figs. 2g,h).

Relative levels of *S1pr1* and *S1pr3* were significantly increased in older mice (16 and 25 weeks of age) whereas levels of *Sphk1* were significantly higher in young mice (2 weeks of age) (Figs. 2i,j; supplemental table 1). The MG from animals at 8 weeks of postnatal development had elevated levels of *S1pr3*, however these cells may have been reactive with significantly elevated levels of *Tgfb2* and *Klf6* (Figs. 2i.j, supplemental table 1), which is characteristic of activated MG that are returning toward a resting phenotype (Hoang et al., 2020; Palazzo et al., 2022).

We next probed patterns of expression of S1P-related genes in scRNA-seq libraries that were established from control retinas and damaged retinas at different times after NMDA-treatment; these libraries were generated and analyzed previously (Campbell et al., 2021a; Campbell et al., 2021b; Hoang et al., 2020; Todd et al., 2019). UMAP plots revealed discrete clusters of all major retinal cell types (Figs. 3a,b). Clusters of cells were labeled based on distinct patterns of gene expression as described in the Methods. Neuronal cells from control and damaged retinas were clustered together regardless of time after NMDA-treatment (Figs. 3a,b). By contrast, resting MG, including cells from 48 and 72 hrs after NMDA, and activated MG from 3, 6, 12, and 24 hrs after NMDA were spatially separated by UMAP embedding (Figs. 3a-c). Activated MG were identified, in part, based on downregulation of genes such as *Slc1a3* and upregulation of genes such as *Vim* or *Nes* (Fig. 3c). We examined patterns of expression and changes in expression levels of S1P-related genes (Figs. 3d-i)*. S1pr4*, *S1pr5*, *Sgpl1*, and *Sphk2* were not detected at significant levels or in very few cells in the retina (not shown). *S1pr2* was expressed by relatively few retinal cells, but *S1pr1*, *S1pr3*, and *Sphk1* were highly expressed by activated MG (Figs. 3d,e). In addition, *S1pr1* was highly expressed by endothelial cells (Fig. 3d). We did not perform detailed analyses of S1P-related genes in endothelial cells because there were less than 350 cells captured in these libraries (Fig. 3b). *S1pr1* and *Sphk1* appeared to be robustly upregulated in activated MG at 3 and 6hrs after NMDA treatment, but downregulated in MG thereafter (Figs. 3d,e). To perform a detailed analysis of S1P related genes in MG, we bioinformatically isolated and re-embedded nearly 6200 MG into a UMAP dimensional reduction (Figs. 3f,g). These MG formed a distinct trajectory of cells with resting MG from control retinas clustered to one side, ++activated MG from 3hrs after NMDA-treated retinas clustered to the other side of the plot, and MG from later times after NMDA bridging the middle of the trajectory (Figs. 3f,g). This analysis revealed that levels of *S1pr1* were relatively low in resting MG, levels were significantly upregulated in ++activated MG at 3hrs after NMDA and further upregulated in +activated MG at 6hrs after NMDA (Figs. 3h,i). Levels of *S1pr1* were significantly downregulated in activated MG and returning to resting MG from retinas at 12-72 hrs after NMDA (Fig. 3i). Levels of *S1pr2* were very low in MG, and levels of *S1pr3* were not significantly changed, but the percentage of MG that expressed *S1pr3* was greatest in MG that were returning to a resting phenotype from retinas 24-72 hrs after NMDA (Figs. 3h,i). By comparison, *Sphk1* was low in resting MG and highly upregulated in ++activated (3hrs after NMDA) and +activated MG (6hrs after NMDA), but significantly downregulated thereafter (Figs. 3h,i). Collectively, these findings suggest that after acute injury levels of *Sphk1* (S1P synthesis) are very rapidly and transiently upregulated, and the upregulation in S1P synthesis is followed by a rapid and transient upregulation in autocrine signaling through *S1pr1* in MG.

**Figure 3:**
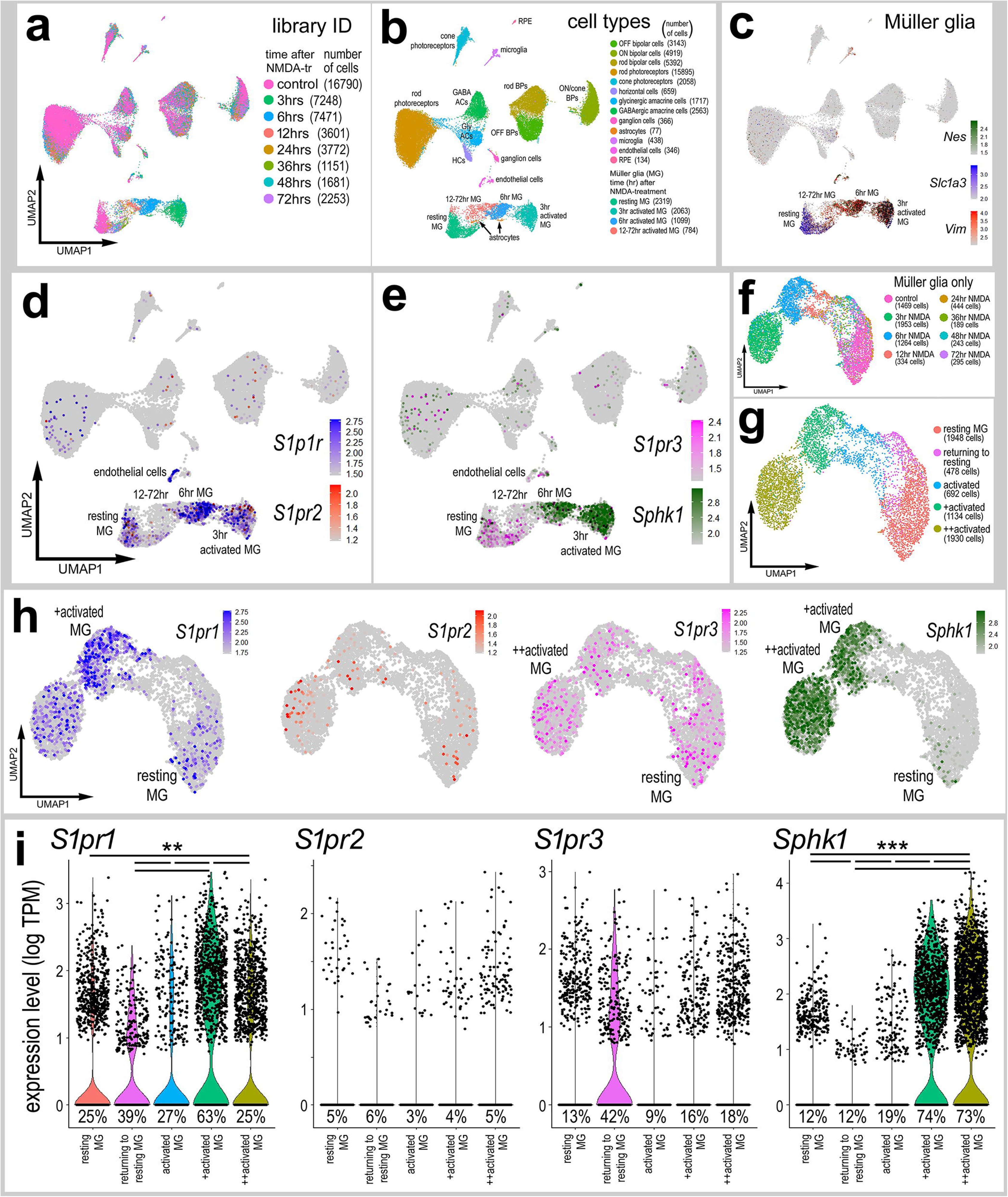
Patterns of expression of S1P-related genes in damaged mouse retinas. scRNA-seq was used to identify patterns of expression of S1P-related factors among retinal cells with the data presented in UMAP (**a-h**) or dot plots (**i**). Aggregate scRNA-seq libraries were generated for cells from control retinas and retinas 3, 6, 12, 24, 36, 48, and 72 hrs after NMDA-treatment. MG were bioinformatically isolated and analyzed from the large aggregate library (**f**-**i**). UMAP-ordered cells formed distinct clusters of neuronal cells, resting MG, activated MG and returning to resting MG based on distinct patterns of gene expression (**c**). UMAP heatmap plots illustrate patterns and levels of expression of S1P receptors *S1pr1, S1pr2, S1pr3,* and S1P metabolism gene *Sphk1* (**h**). Dot plots illustrate relative levels of expression (heatmap) and percent expression (dot size) in MG, activated MG, and returning to resting MG (**i**). Significance of difference was determined by using a Wilcox rank sum with Bonferoni correction (supplemental table 1). Abbreviations: MG, Müller glia; NMDA, N-methyl-D-aspartate; UMAP, uniform manifold approximation and projection.

### Validation of patterns of expression of *S1pr1* and *Sphk1*

To validate findings from scRNA-seq libraries, we applied different immunolabeling strategies using different antibodies to S1pr1 and Sphk1 (Table 1). However, none of these antibodies produced plausible patterns of labeling regardless of different antigen retrieval strategies including weak fixation, methanol washes, sodium citrate washes, exclusion of detergent, or applying different types of detergents (not shown). Thus, we applied fluorescent *in situ* hybridization (FISH) strategies with probes for *S1pr1* and *Sphk1*. We did not probe for *S1pr2* or *S1pr3* because the scRNA-data indicated very low levels of expression in the retina. In undamaged retinas, *S1pr1* puncta were associated with Sox2-labeled MG nuclei (Fig. 4a). *S1pr1* FISH signal was also detected in putative endothelial cells associated with auto-fluorescent red blood cells that were scattered across retina (Fig. 4a). Consistent with findings from scRNA-seq, we found an increase in *S1pr1* puncta associated with Sox2-labeled MG nuclei at 4 hrs after NMDA-treatment (Fig. 4a).

**Figure 4:**
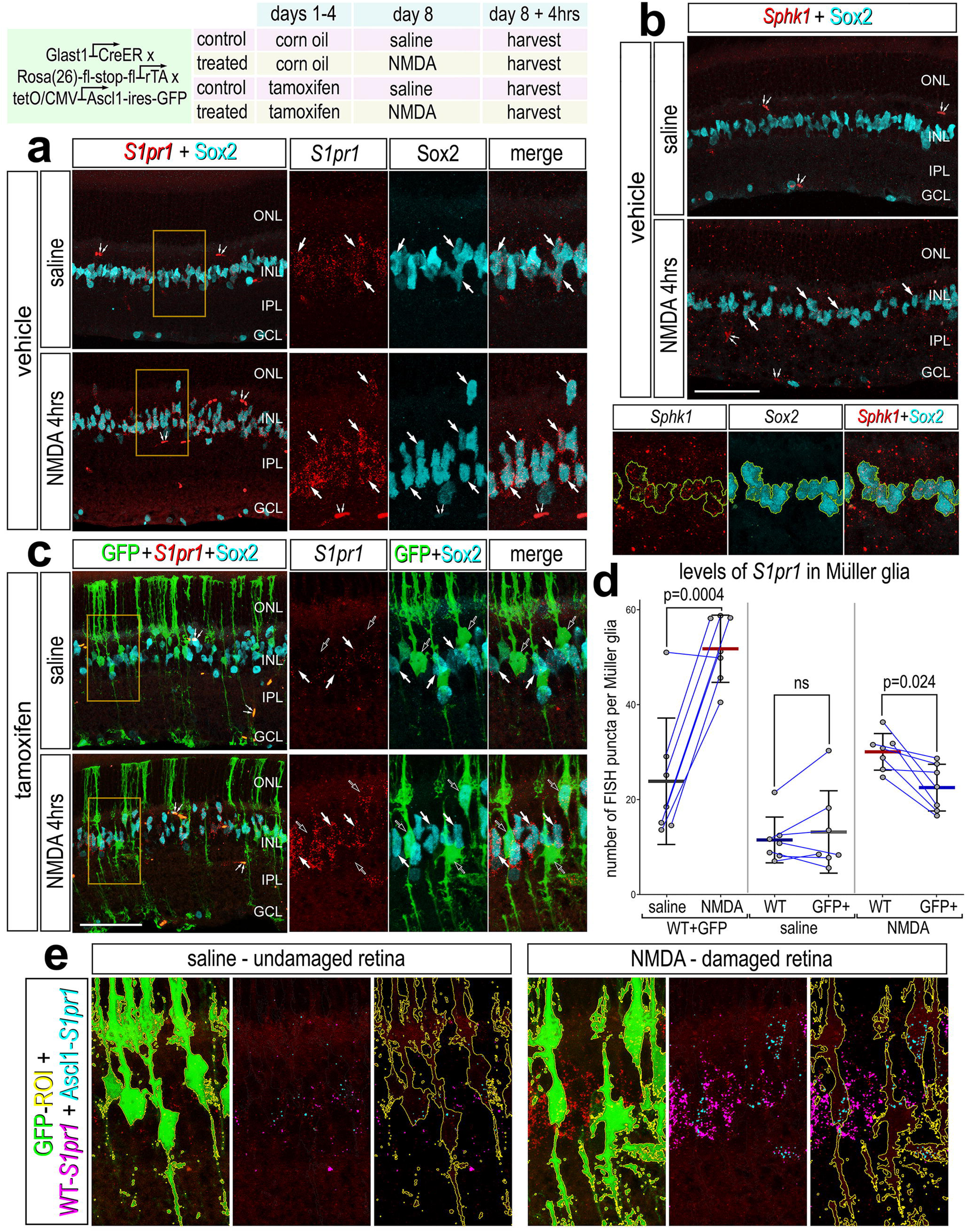
Fluorescence *in situ* hybridization (FISH) for *S1PR1* and *SPHK1*. *Glast-CreER:LNL-tTA:tetO-mAscl1-ires-GFP* mice received IP injections of tamoxifen or corn oil once daily for 4 consecutive days. Eyes received intravitreal injections with NMDA or saline, and retinas were collected 4 hrs after NMDA treatment. Retinal sections were labeled with antibodies to Sox2 (cyan), GFP (green) or FISH probes to *S1PR1* (red puncta; **a**, **c**) or *SPHK1* (red puncta; **b**). Regions of interest (yellow boxes) are enlarged 1.8-fold and displayed in adjacent channels. Solid arrows indicate Sox2^+^/GFP^low^ MG nuclei, and hollow arrows indicate Sox2^+^/GFP^high^ MG nuclei. The area occupied by Sox2^+^ (**b**) or GFP^+^ cells (**e**) was selected to distinguish FISH puncta labeling within these regions. Small double arrows indicate endothelial cells. Calibration bars panels **a**, **b**, **c**, and **e** represent 50 µm. Histograms represent the mean (bar ± SD) number of FISH puncta per MG and each dot represents one biological replicate (**d**). Significance of difference (p-values) was determined by using a paired t-test. Abbreviations: ONL, outer nuclear layer; INL, inner nuclear layer; IPL, inner plexiform layer; GCL, ganglion cell layer; NMDA, N-methyl-D-aspartate; ns, not significant.

In undamaged retinas, there were very few *Sphk1* FISH puncta scattered randomly across the retina (Fig. 4b). At 4hrs after NMDA-treatment there were numerous *Sphk1* puncta scattered across the retina and some of these puncta were associated with Sox2^+^ MG nuclei (Fig. 4b). According to scRNA-seq data, levels of *Sphk1* are increased in MG at 3-6 hrs after NMDA, whereas the scattered distribution of *Sphk1* FISH puncta did not reveal MG-specific upregulation of transcripts.

To investigate whether S1P-signaling through *S1pr1* is altered during the reprogramming of MG into progenitor-like cells, we applied FISH probes to retinas where the MG have been forced to express the neurogenic basic Helix-Loop-Helix (bHLH) transcription factor *Ascl1 (Glast-CreER:LNL-tTA:tetO-mAscl1-ires-GFP)* (Jorstad et al., 2017). In adult mouse retinas *in vivo*, inducible forced expression of *Ascl1* in MG, combined with NMDA-induced damage and HDAC inhibitor (trichostatin A), stimulates the reprogramming of MG into bipolar-like cells that integrate into circuitry and respond to visual stimuli (Jorstad et al., 2017). Ascl1 over-expressing (Ascl1-OE) MG were identified by GFP expression which is linked to the *Ascl1* transgene via an internal ribosomal entry sequence (IRES) (Fig. 4). Counts were made for *S1pr1* FISH puncta associated with Ascl1-OE MG (Sox2 ^+^/GFP^+^ cells) and for puncta associated with neighboring WT MG (Sox2^+^/GFP^-^ cells) within in the same section. In undamaged retinas, numbers of *S1pr1* puncta in Ascl1-OE MG were not significantly different from WT MG (Figs. 4c-e). In damaged retinas, there were significantly more *S1pr1* FISH puncta per MG in NMDA-damaged retinas compared to numbers seen in saline-treated retinas (Figs. 4c-e), consistent with scRNA-seq data. In damaged retinas, with Ascl1-OE MG, numbers of *S1pr1* FISH puncta were significantly reduced in Ascl1-OE MG compared to numbers seen in WT MG (Figs. 4c-e). Collectively, these findings indicate that *S1pr1* and *Sphk1* transcripts are upregulated in response to NMDA damage but downregulated in response to forced expression of *Ascl1* in resting and damaged retinas.

### S1P-signaling and reprogramming of Ascl1-over expressing MG

To investigate the role of S1P-signaling in Ascl1-mediated neurogenesis, we applied small molecule inhibitors to Sphk1 (PF543) and S1pr1/3 (VPC23019) to the retinas of *Ascl1-OE mice.* Retinas were treated with NMDA and inhibitors, inhibitors at 1 day post-injury (DPI), TSA at 2 DPI, and harvested at 16 DPI (Fig. 5). Application of PF543 or VPC23019 alone suppressed glial differentiation but did not significantly influence numbers of differentiating neurons (not shown). Application of Sphk1+S1pr1/3 inhibitors significantly decreased the percent of GFP^+^/Sox2^+^ cells (Figs. 5a-c), indicating that the S1P pathway promotes glial dedifferentiation in Ascl1-OE MG. By comparison, application of Sphk1+S1pr1/3 inhibitors significantly increased the percent of GFP^+^/Otx2^+^ cells (Figs. 5d-f), indicating that S1P pathway suppresses neuronal dedifferentiation in Ascl1-OE MG. Sphk1+S1pr1/3 inhibitors had no significant effect upon total numbers of proliferating EdU-positive cells or numbers of EdU^+^/GFP^+^ cells (Fig. 5g). Collectively, these findings indicate that inhibition of S1pr1/3 and Sphk1 in *Ascl1*-OE MG enhances the differentiation of bipolar-like cells, while glial phenotype was suppressed.

**Figure 5:**
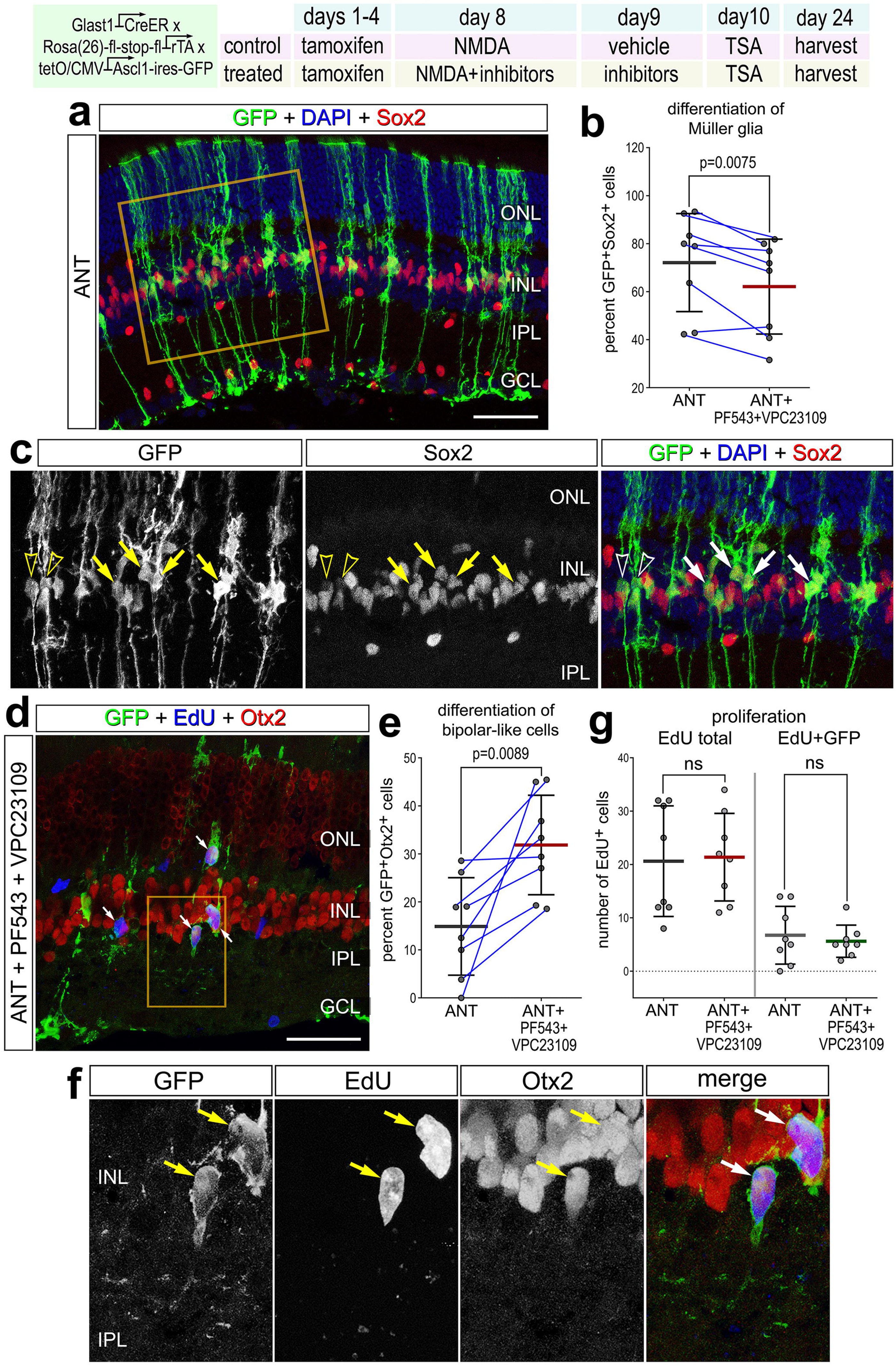
Inhibition of S1P-signaling enhances Ascl1-mediated reprogramming of MG into neurogenic progenitor cells. *Glast-CreER:LNL-tTA:tetO-mAscl1-ires-GFP* mice were IP injected with tamoxifen or corn oil once daily for 4 days. Eyes were intravitreally injected NMDA and either a vehicle control or inhibitors to Sphk1 (PF543) and S1pr1/3 (VPC23019). At 24HPI, eyes were treated with vehicle or inhibitors. Eyes were treated with TSA at 48HPI. Retinas were collected 24 days from the first tamoxifen injection. Retinal sections were labeled with DAPI (**a**, **c**) or EdU (**d**, **f**) and antibodies to GFP (green) and Sox2 (red, **a**, **c**) or Otx2 (red, **d**, **f**). Regions of interest (yellow boxes) are enlarged 1.5 (**a,c**) or 2-fold (**d,f)** and displayed in separate panels. Solid arrows indicate GFP^+^/Sox2^high^ cells in **a** and **c**, and hollow arrowheads indicate GFP^+^/Sox2^low^ cells. In **d** and **f**, solid arrows indicate GFP^+^/Otx2^+^/EdU^+^ cells. Calibration bars in panels **a**, **b**, **c**, and **e** represent 50 µm. Histograms represent the mean (bar ± SD) where each dot represents one biological replicate, and blue lines connect control and treated eyes from the same individual (**b**, **e**, **g**). Significance of difference (p-values) was determined by using a paired t-test. Abbreviations: ONL, outer nuclear layer; INL, inner nuclear layer; IPL, inner plexiform layer; GCL, ganglion cell layer; NMDA, N-methyl-D-aspartate; ns, not significant.

### S1P/S1pr1-signaling activates different second messenger pathways in MG

S1P receptors differ by their affinity to S1P and by their coupling to different G-proteins, thereby resulting in distinct cellular responses and physiological outcomes. S1pr1 has been shown to predominantly couple to Gi, activating Phospholipase C (PLC), MAPK/Erk, and PI3K/AKT pathways (Pyne and Pyne, 2017), and secondarily impact NFκB-signaling (reviewed by Sun et al., 2024). Thus, we investigated whether pharmacological activation or inhibition of S1P-related receptors and enzymes influenced ERK phosphorylation, NFκB activation, mTor activation (pS6) and cFos expression in normal and damaged retinas.

Eyes were treated with a single intraocular injection of S1P or S1pr1 agonists (CYM5422 or SEW2871) and harvested retinas 24 hours later. Alternatively, eyes were injected with an S1pr1 agonist (SEW2871), S1pr1 antagonist (MT1303), S1pr3 antagonist (TY52156), S1pr modulator (FTY720), Sphk1 antagonist (PF543), or S1P lyase antagonist (S1PL-in-31) before and with NMDA, and retinas were harvested 24 hours after the last injection. See Figure 1 for the different sites of action for these drugs. FTY720, or Fingolimod, is a clinically approved S1pr1-modulator which initially acts as an agonist to S1pr1 but later induces receptor internalization and degradation (Chun et al., 2021).

In undamaged retinas treated with exogenous S1P or S1PR1 activator, we applied antibodies to different second messengers for cell signaling pathways in the retina (Figs. 6a-d). In retinas treated with S1pr1 agonist (SEW2871), pERK1/2 levels were significantly elevated in Sox2-expressing MG (Figs. 6a,b). However, intravitreal delivery of S1P did not result in the accumulation of pERK1/2 in MG (Fig. 6b). Levels of immediate early gene cFos or mTor target pS6 were not significantly different between treatment groups (Figs. 6c,d), indicating that these pathways are not affected by S1P-signaling. We have recently reported that NFκB-signaling is rapidly, transiently and selectively activated in MG in damaged mouse retinas, and this activation is mediated by pro-inflammatory cytokines produced by reactive microglia (Palazzo et al., 2022; Palazzo et al., 2023). Since S1P-signaling is known to activate inflammatory responses via NFκB in many cell types, including endothelial cells, cancer cells and macrophages (Campos et al., 2016; Del Gaudio et al., 2020; Liang et al., 2013; Xiao et al., 2018; Zheng et al., 2019), we sought to determine whether S1P influences NFκB activation in MG in normal and damaged retinas. We used NFκB reporter mice wherein eGFP expression is driven by a chimeric promoter containing three HIV-NFκB cis-regulatory elements (Magness et al., 2004). In the NFκB-reporter mice there is MG-specific NFκB activation in retinas treated with NMDA, cytokines, and different growth factors (Palazzo et al., 2023) (Fig. 6e). In undamaged retinas, we found that S1pr1 activation, via S1P or different agonists, significantly increased numbers of Sox2-positive MG that express NFκB reporter in undamaged retinas (Figs. 6f,g). Additionally, S1pr1 agonist treatment increased numbers of NFκB-expressing MG in damaged retinas (Fig 6i). Consistent with findings from studies using S1pr1 agonist, numbers of NFκB-GFP-expressing MG were increased in retinas treated with S1P lyase inhibitor, decreased in retinas treated with S1pr1 antagonist, decreased in retinas treated with Sphk1 inhibitor (Figs. 6h,i), whereas S1pr3 antagonist (Fig. 6i) and S1pr1-modulator (not shown) had no significant effects on numbers of MG that expressed NFκB reporter in damaged retinas. Collectively, these findings suggest that autocrine S1P-signaling via Sphk1-mediated synthesis of S1P and activation of S1pr1 activates NFκB-signaling in MG in normal and damaged retinas.

**Figure 6:**
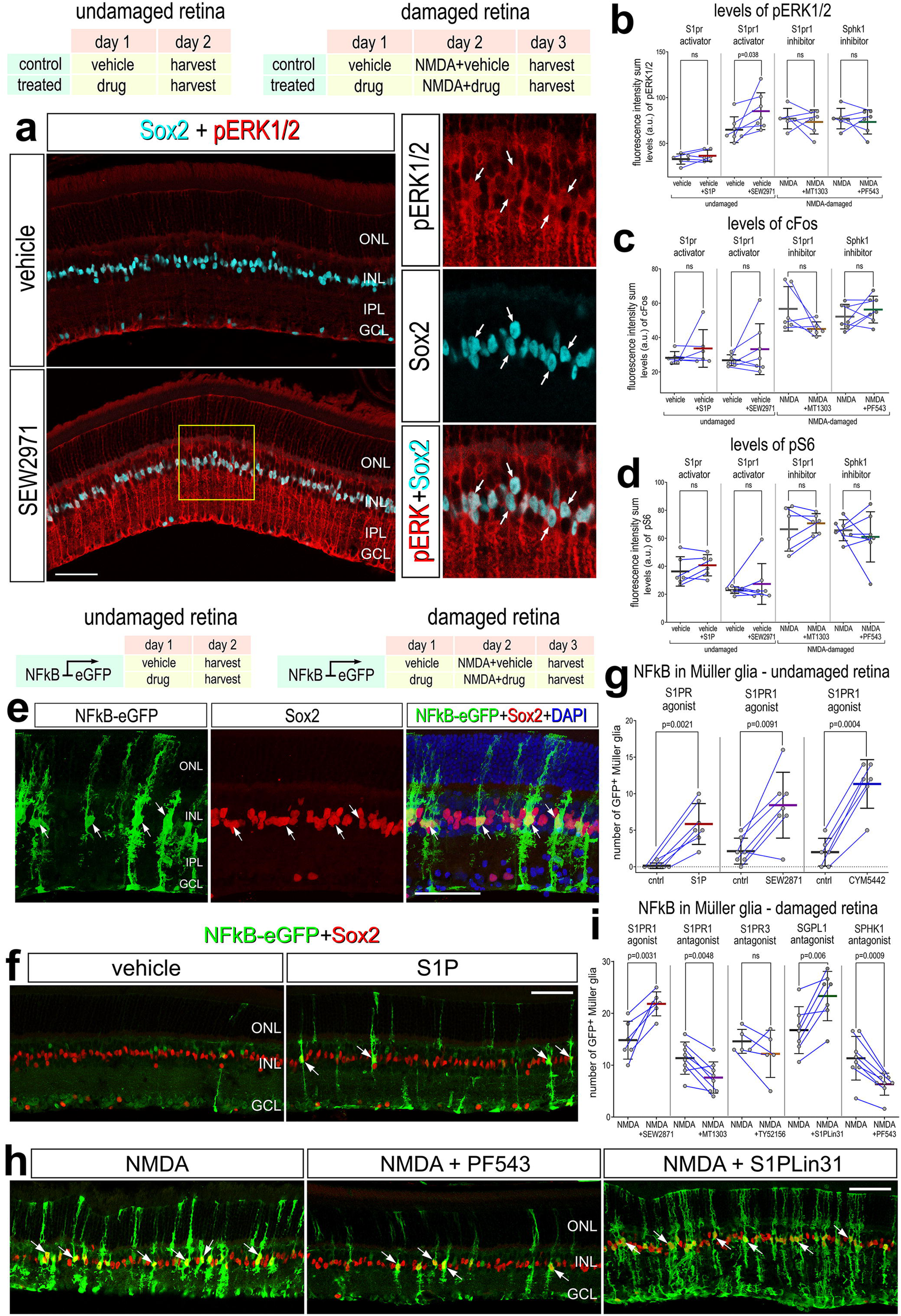
Activation of cell signaling pathways by S1P, agonists and antagonists. Retinas were obtained from undamaged or NMDA-injured eyes treated with a vehicle control or a small molecule agonist/antagonist. Retinal sections were labeled with DAPI (**e**) and antibodies to Sox2 (cyan, **a**; red, **e**, **f**, **h**), pERK1/2 (red, **a**), cFos, pS6, or NFκB - GFP (**e**, **f**, **h**). Arrows indicate the nuclei of MG. Calibration bars in panels **a**, **e**, **f**, and **h** represent 50 µm. Regions of interest (yellow boxes) are enlarged 1.8-fold and displayed in separate panels. Histograms in **b**, **c**, **d**, **g**, **i** illustrate the mean (bar ± SD) fluorescence intensity sum for pERK1/2, cFos, pS6 or NFκB-eGFP in MG, where each dot represents one biological replicate, and blue lines connect replicates from control and treated retinas from one individual. Significance of difference (p-values) was determined by using a paired t-test. Abbreviations: ONL, outer nuclear layer; INL, inner nuclear layer; IPL, inner plexiform layer; EdU, 5-Ethynyl-2′-deoxyuridine; NMDA, N-methyl-D-aspartate; ns, not significant.

### NFκB-signaling regulates the expression of S1P-related genes in MG

S1P-signaling is known to activate NFκB ((Alvarez et al., 2010) and Fig.5), and NFκB is known to reciprocally regulate S1P-signaling (Hutami et al., 2017; O’Sullivan et al., 2014). Accordingly, we probed for levels of S1P-related genes in scRNA-seq of retinas where NFκB-signaling has been selectively deleted from MG. We analyzed libraries that we generated and described in a prior study (Palazzo et al., 2022). Rlbp1-CreERT mice were crossed to mice carrying floxed alleles of *Ikkb* to prevent NFκB-signaling in MG. Activation of canonical NFκB-signaling involves the IKK complex (Ikka and Ikkb) which is activated to phosphorylate IkBα/IkBβ, thereby releasing NFκB transcription factors (p65 and p50) and allowing them to translocate into the nucleus to regulate transcription of target genes (Zhang et al., 2017). Deletion of *Ikkb* blocks signaling through the NFκB pathway by preventing IKK-mediated phosphorylation and degradation of IkBa/IkBb, thereby causing sequestration of NFκB transcription factors in the cytoplasm (Li et al., 2003; Zhang et al., 2017, 30).

Consistent with patterns of expression seen in scRNA-seq libraries in WT retinas (Fig. 2), at 8hrs after NMDA-treatment *S1pr1*, *S1pr3* and *Sphk1* were highly expressed by MG (Figs. 7a-f). In addition, *Sgpl1* was detected in immune cells and MG (Figs. 7a-f), consistent with patterns of expression seen in WT retinas. We bioinformatically isolated the nearly 1400 MG to permit a more detailed analysis. In MG where *Ikkb* was deleted, levels of *S1pr1* were significantly upregulated, whereas levels of *S1pr3* and *Sphk1* were significantly reduced, and levels of *Sgpl1* were unchanged (Fig. 7h). We next sought to validate changes in patterns of *S1pr1* expression in retinas where NFκB-signaling was inhibited. We applied a small molecule NFκB inhibitor, PGJ2, which has been shown to potently block NFκB reporter activation in NMDA-damaged mouse retinas (Palazzo et al., 2022), followed by FISH for *S1pr1*. We found that the number of S1pr1 FISH puncta per MG were significantly increased in NMDA-damaged retinas treated with PGJ2 (Figs. 7i,j). Collectively, these findings suggest that activation of S1pr1/3 and downstream NFκB-signaling reciprocally coordinate expression levels in MG in damaged retinas.

**Figure 7:**
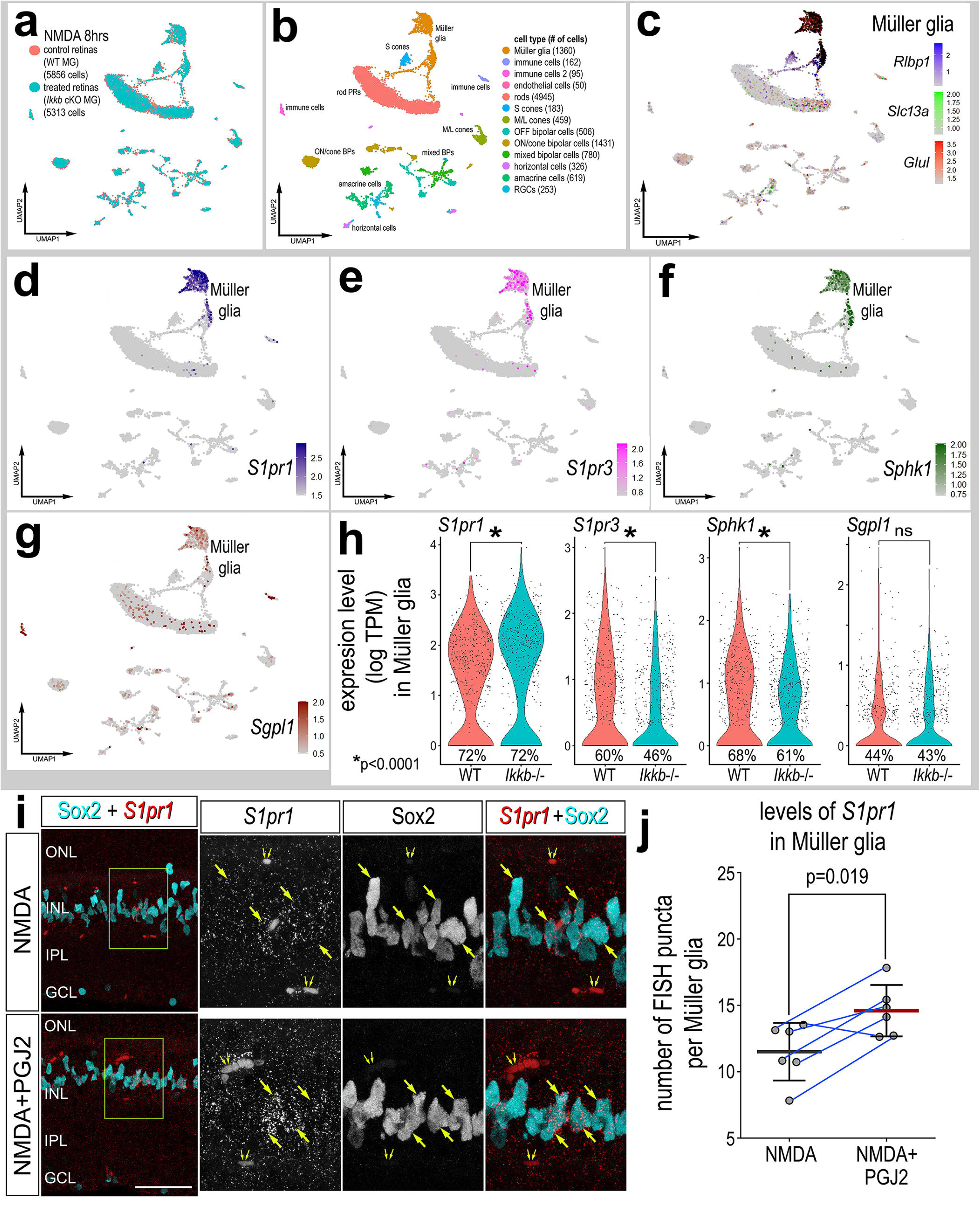
NFκB-signaling activity regulates S1P-related gene expression. scRNA-seq was used to identify patterns of expression of S1P-related factors among retinal cells with the data presented in UMAP (**a-g**) or violin plots (**h**). scRNA-seq libraries were generated for cells from Rlbp1-creERT (Ikkb-WT) and Rlbp1-creERT:Ikkbfl/fl (Ikkb-cKO) retinas 8 h after NMDA (**a**). UMAP-ordered cells formed distinct clusters of neuronal cells, MG, and immune cells based on distinct patterns of gene expression (**b**, **c**). UMAP heatmap plots illustrate patterns and levels of expression of S1P receptors *S1pr1* and *S1pr3* (**d**, **e**), and S1P metabolism genes *Sphk1* and *Sgpl1,* and *SPNS2* (**f, g**) across the whole retina. Violin plots illustrate relative levels of expression (heatmap) and percent expression (dot size) in WT MG and Ikkb-cKO MG (**h**). WT mouse eyes were intravitreally injected with NMDA and vehicle control or NFκB inhibitor PGJ2, and retinas were collected 4 h after NMDA. Retinal sections were labeled with antibodies to Sox2 (cyan) and a FISH probe to *S1pr1* (red puncta; **i**). Regions of interest (yellow boxes) are enlarged 2-fold and displayed in adjacent channels. Solid arrows indicate Sox2+ nuclei, whereas small double arrows indicate RFP+ endothelial cells. Calibration bar in panel **i** represents 50 µm. Histogram represents the mean (bar ± SD) where each dot represents one biological replicate and blue lines connect control and treated eyes from the same individual (**j**). Significance of difference (p-values) was determined by using a Wilcox rank sum with Bonferoni correction (**h**) or by using paired t-test (**j**). Abbreviations: ONL, outer nuclear layer; INL, inner nuclear layer; IPL, inner plexiform layer; GCL, ganglion cell layer; NMDA, N-methyl-D-aspartate; ns, not significant; UMAP, uniform manifold approximation and projection.

### The accumulation of immune cells in damaged retinas is influenced by *S1pr1*, but not *Sphk1*

To further investigate how signaling through S1pr1 and synthesis of S1P via Sphk1 in MG influences the retina we generated conditional knock-out (cKO) mice. We crossed Rlbp1-CreERT mice with mice carrying floxed alleles of *S1pr1* or *Sphk1*. Mice were treated with vehicle or tamoxifen for 4 consecutive days, followed by 3 days to allow recombination and tamoxifen to clear. Eyes were injected with saline or NMDA, and retinas harvested 2 days after NMDA (Fig. 8). We probed for the accumulation of resident microglia and peripheral immune cells that are known to migrate into the retina after injury (White et al., 2017b; Yu et al., 2020). Resident microglia and macrophages are CD45-low, whereas peripheral monocyte derived macrophages are CD45-high expressing cells (O’Koren et al., 2016; Sedgwick et al., 1991). Accordingly, we identified CD45+/Iba1-cells as putative infiltrating monocyte-derived macrophages and CD45+/Iba+ cells as microglia. We found that cKO of *S1pr1* or *Sphk1* had no effect on numbers of microglia or peripheral monocytes in undamaged retinas (Figs. 8a-d). In WT retinas, two days after NMDA-treatment we found significant increases in numbers of microglia, but not peripheral monocytes (Figs. 8a-d). Numbers of microglia and peripheral monocytes in NMDA-damaged retinas with *S1pr1*-/- MG were significantly increased when compared to numbers seen in undamaged retinas with *S1pr1*-/- MG and NMDA-damaged retinas with WT MG (Figs. 8a,b). By comparison, there was no significant difference in the number of peripheral monocytes NMDA-damaged retinas with *Sphk1*-/- MG compared to numbers seen in NMDA-damaged retinas with WT MG (Figs. 8a-c). However, numbers of microglia in damaged retinas with *Sphk1*-/- MG were significantly increased compared to numbers seen in undamaged retinas with *Sphk1*-/- MG (Figs. 8a,b,d). In summary, the deletion of *S1pr1*, but not *Sphk1*, from MG causes the increased accumulation of peripheral monocytes and microglia in acutely damaged retinas.

**Figure 8:**
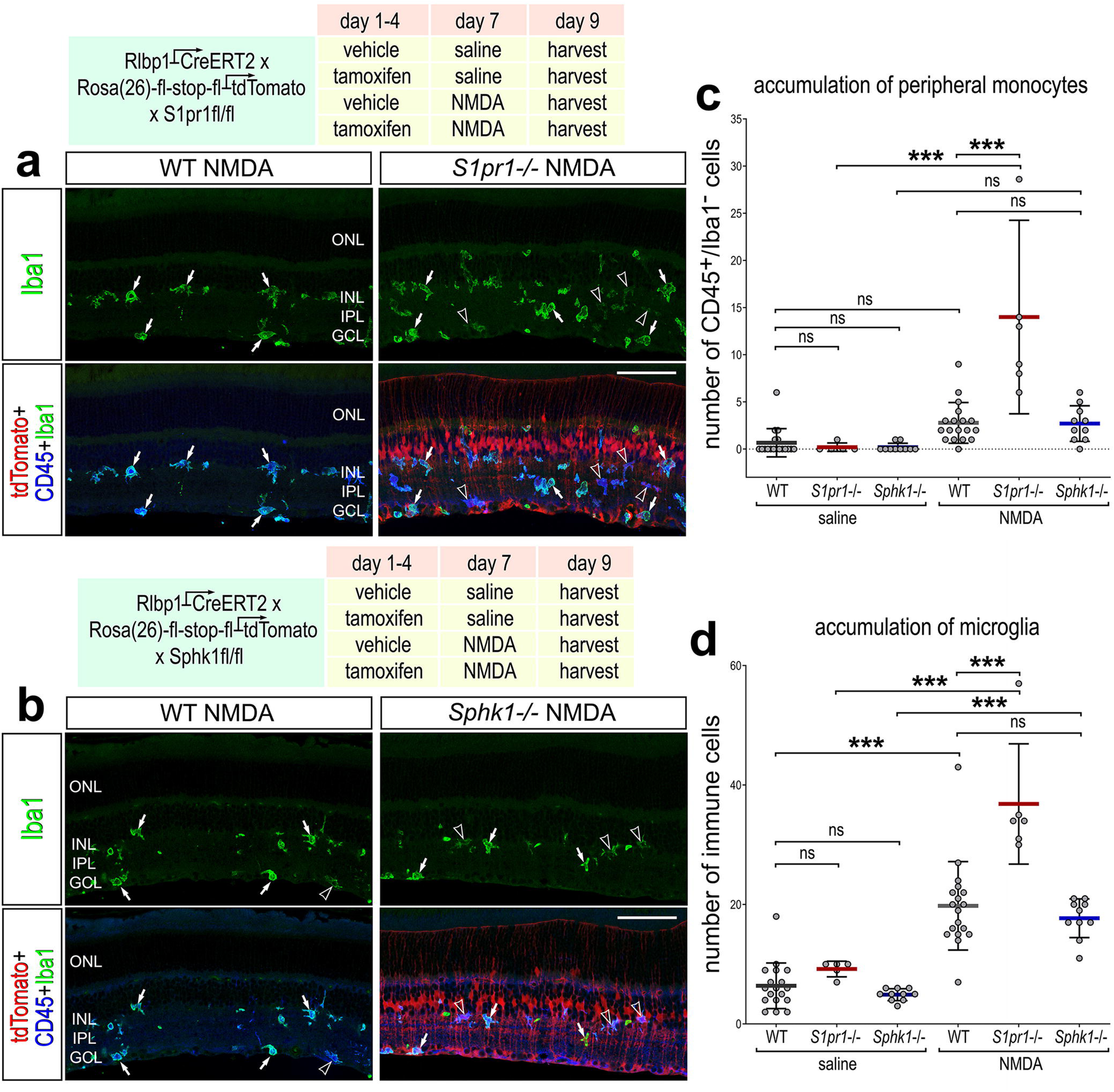
Immune cell accumulation in S1pr1 KO and Sphk1 KO retinas. *Rlbp1-CreERT:S1pr1^fl/fl^* and *Rlbp1-CreERT:Sphk1^fl/fl^* mice were IP injected with tamoxifen or corn oil once daily for 4 days. Eyes were intravitreally injected withNMDA or saline control, and retinas were collected 48HPI. Retinal sections were labeled with antibodies to Iba1 (blue) and CD45 (green) (**a**, **b**). Solid arrows indicate Iba1^high^/CD45^high^ cells, and hollow arrowheads indicate Iba1^high^/CD45^low^ cells. Calibration bars in panels **a** and **b** represent 50 µm. Histograms represent the mean (bar ± SD) where each dot represents one biological replicate (**c**, **d**). Significance of difference (p-values) was determined by using ANOVA with Sidak correction. Abbreviations: ONL, outer nuclear layer; INL, inner nuclear layer; IPL, inner plexiform layer; GCL, ganglion cell layer; NMDA, N-methyl-D-aspartate; ns, not significant.

### S1P-signaling in MG regulates neuronal survival in damaged retinas

NMDA-treatment is known to selectively destroy bipolar cells, amacrine cells and retinal ganglion cells, but not MG in the mouse retina (Todd et al., 2019). Since neuroinflammation is known to impact neuronal survival in NMDA-damaged retinas (Todd et al., 2019), and S1P-signaling influences the NFκB pathway and the accumulation of microglia/macrophage (current study), we investigated whether S1P influences cell death and neuronal. In undamaged retinas with WT or *S1pr1* cKO MG we found no evidence for dying TUNEL-positive cells (Figs. 9a-c). We found significant increases in numbers TUNEL-positive cells in the INL/ONL or GCL in NMDA retinas at 2 days after treatment, and these numbers were significantly reduced in the GCL with MG-specific cKO of *S1pr1* (Figs. 9a-c). Consistent with these findings, we found significantly more surviving retinal ganglion cells (RGCs) labeled for Brn3a or HuC/D in the GCL at 2 weeks after NMDA-treatment in *S1pr1* cKO retinas compared to numbers seen in WT retinas (Figs. 9d-g). However, we did not find a significant difference in numbers of HuC/D+ amacrine cells in the INL at 2 weeks after NMDA-treatment in retinas with WT MG and retinas with *S1pr1-/-* MG (Figs. 9f,h).

**Figure 9:**
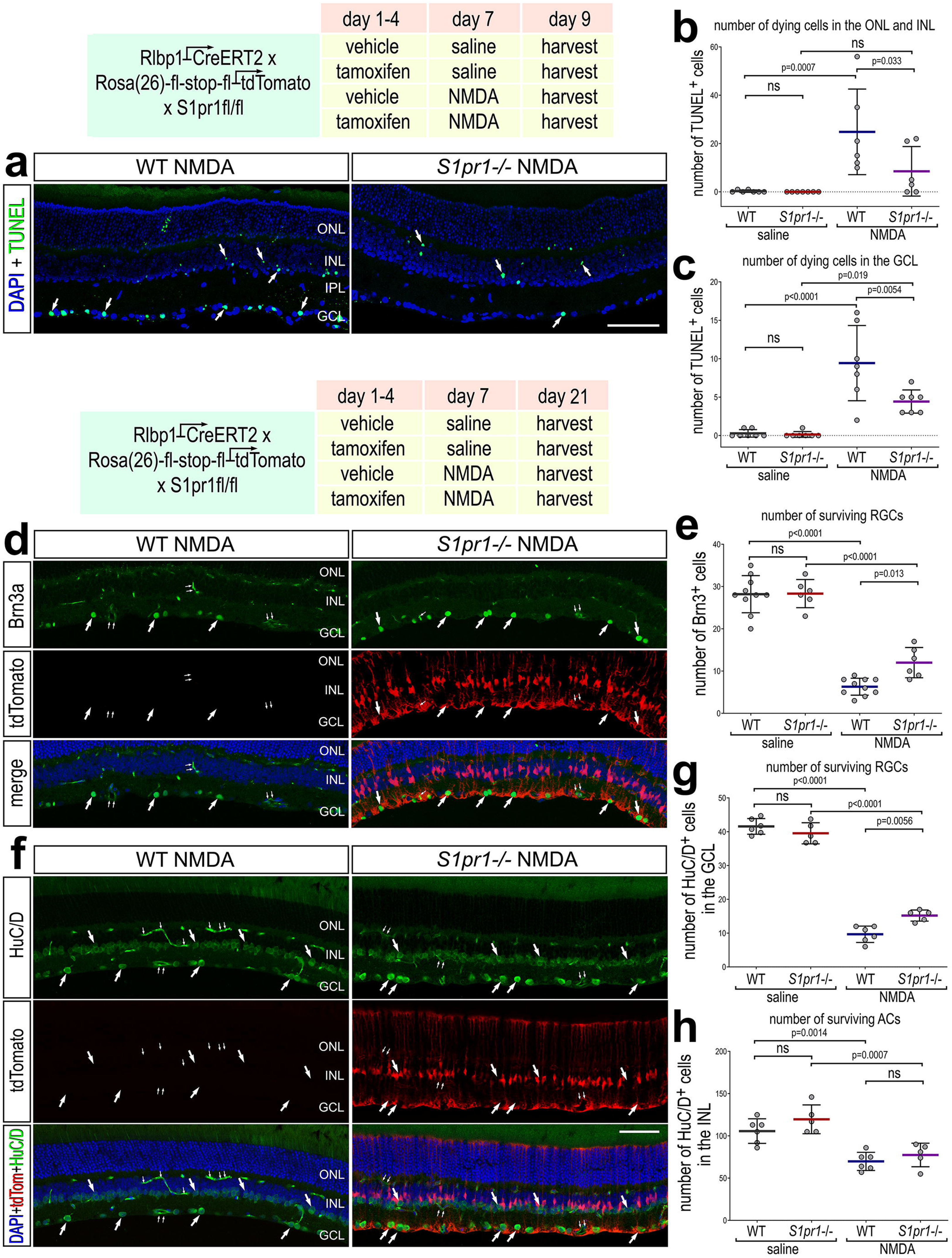
cKO of *S1pr1* from MG is neuroprotective to retinal ganglion cells. Sections of the retina were labeled for fragmented DNA (TUNEL; green; **a**) and DAPI (blue), or Brn3a (green; **d**), tdTomato (red) and DAPI (blue), or HuC/D (green; **f**), tdTomato (red) and DAPI (blue; **f**). Arrows indicate TUNEL+ nuclei (**a**) or Brn3a+ nuclei (**d**) or HuC/D+ cells (**f**) and small double arrows indicate GFP+ endothelial cells. Calibration bars in panels **a**, **d**, and **f**, represent 50 µm. Histograms represent the mean (bar ± SD) where each dot represents one biological replicate (**b**, **c**, **e**, **g**, **h**). Significance of difference (p-values) was determined by using ANOVA with Sidak correction. Abbreviations: ONL, outer nuclear layer; INL, inner nuclear layer; IPL, inner plexiform layer; GCL, ganglion cell layer; NMDA, N-methyl-D-aspartate; ns, not significant; TUNEL, terminal deoxynucleotidyl transferase dUTP nick end labeling.

In retinas with cKO of *Sphk1* in MG, we found significantly fewer TUNEL+ cells in the INL and ONL compared to WT retinas (Figs. 10a-c). We did not find a significant difference in numbers of RGCs labeled for Brn3a or HuC/D at 2 weeks after NMDA-treatment in retinas with *Sphk1-/-* MG compared to numbers seen in retinas with WT MG (Figs. 10d-e). However, we found a significant increase in numbers of HuC/D+ or Pax6+ amacrine cells in the INL at 2 weeks after NMDA-treatment in retinas with *Sphk1-/-* MG (Figs. 10f-i). Collectively, these findings indicate that cKO or *S1pr1* or *Sphk1* in MG is neuroprotective to inner retinal neurons that are damaged by NMDA. We next investigated whether pharmacological inhibition of S1pr1, Sphk1 or Sgpl1 influences cell death. Eyes were injected with vehicle, MT1303 (S1pr1 inhibitor), PF543 (Sphk1 inhibitor), FTY720 (S1pr1 modulator) or S1PLin31 (SGPL1 inhibitor) before and with NMDA, and retinas harvested 1 day after NMDA-treatment. We observed significantly reduced numbers of TUNEL+ cells in damaged retinas treated with Sphk1 inhibitor and S1pr1 modulator (Figs. 11a,b). In retinas treated with Sphk1 inhibitor, numbers of TUNEL+ cell were reduced in the INL/ONL and GCL, whereas in FTY710-treated retinas cell death was reduced only in the GCL (Figs. 11a,c,d). Complementary to these findings, treatment of retinas with S1PLin31 resulted in significant increases in numbers of TUNEL+ cells in the INL (Figs. 11a-d). In damaged retinas treated with S1pr1 antagonist numbers of TUNEL+ cells were not significantly different from control retinas (Figs. 11a-d). Collectively, these findings suggest inhibition of S1P synthesis suppresses and inhibition of S1P degradation increases cell death in acutely damaged retinas, and these effects are not mediated by diminished signaling through S1pr1.

**Figure 10:**
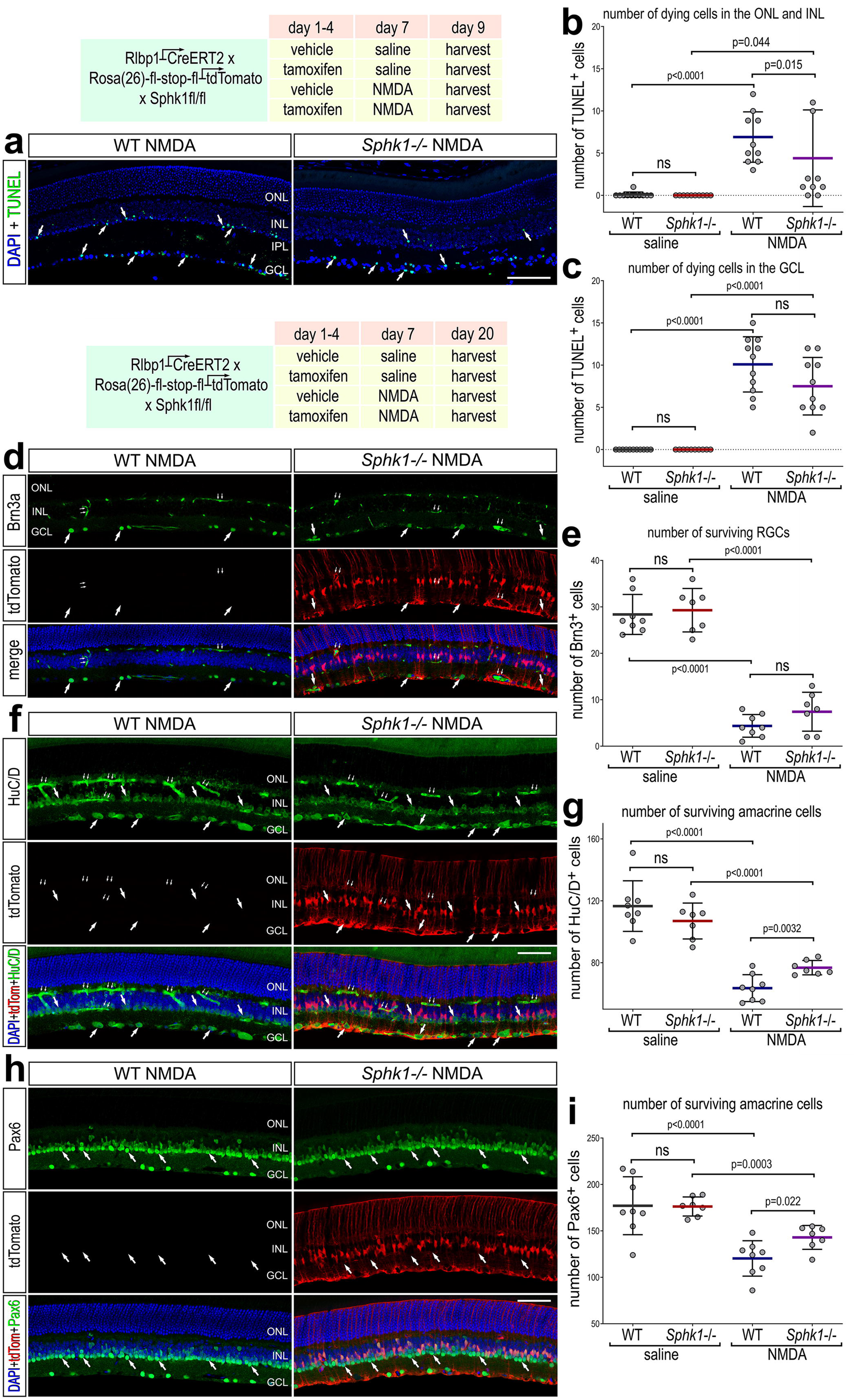
cKO of *Sphk1* from MG is neuroprotective to inner retinal neurons. Sections of the retina were labeled for fragmented DNA (TUNEL; green; **a**) and DAPI (blue), or Brn3a (green; **d**), tdTomato (red) and DAPI (blue), or HuC/D (green; **f**), tdTomato (red) and DAPI (blue), or Pax6 (green; **h**), tdTomato (red) and DAPI (blue). Single arrows indicate TUNEL+ nuclei (**a**) or Brn3a+ nuclei (**d**) or HuC/D+ cells (**f**), and small double arrows indicate GFP+ endothelial cells. Calibration bars in panels **a**, **d**, **f**, and **h** represent 50 µm. Histograms represent the mean (bar ± SD) where each dot represents one biological replicate (**b**, **c**, **e**, **g**, **i**). Significance of difference (p-values) was determined by using ANOVA with Sidak correction. ONL, outer nuclear layer; INL, inner nuclear layer; IPL, inner plexiform layer; GCL, ganglion cell layer; NMDA, N-methyl-D-aspartate; ns, not significant; TUNEL, terminal deoxynucleotidyl transferase dUTP nick end labeling.

**Figure 11:**
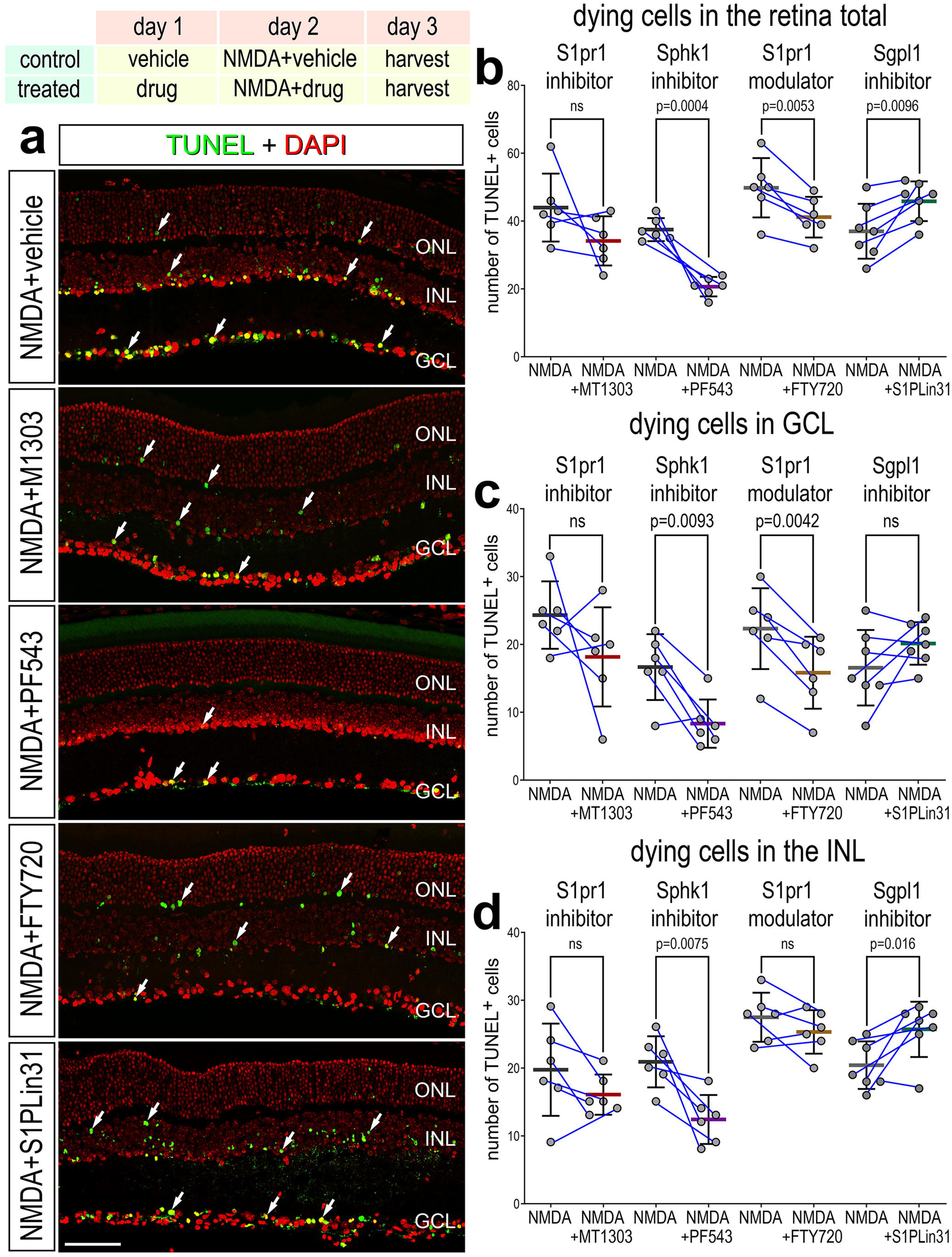
Pharmacological activation or inhibition of S1P-signaling influences cell death in damaged retinas. Retinas were obtained from NMDA-injured eyes treated with a vehicle control or a small molecule agonist/antagonist. Sections of the retina were labeled for fragmented DNA (TUNEL; green) and DAPI (red; **a**). Arrows indicate TUNEL+ puncta. Calibration bar in panel a represents 50 µm. Histograms illustrate mean (bar ± SD) numbers of TUNEL+ cells in the whole retina (**b**), GCL (**c**), and INL (**d**), Each dot represents one biological replicate, and blue lines connect replicates from control and treated retinas from one individual. Significance of difference (p-values) was determined by using a paired t-test. Abbreviations: ONL, outer nuclear layer; INL, inner nuclear layer; IPL, inner plexiform layer; GCL, ganglion cell layer; NMDA, N-methyl-D-aspartate; ns, not significant; TUNEL, terminal deoxynucleotidyl transferase dUTP nick end labeling.

Todd and colleagues reported that microglia suppress Ascl1-mediated neurogenesis without influencing cell survival in mouse retinas (Todd et al., 2020). Additionally, we have previously reported that NFκB-signaling in MG is induced by Il1α, Il1β and TNF from reactive microglia, and this signaling regulates genes responsible for neuron survival (Palazzo et al., 2022; Palazzo et al., 2023; Todd et al., 2019). Accordingly, we investigated whether the neuroprotective effects of S1pr1 inhibition are dependent on immune cell proliferation and recruitment. We ablated microglia in *S1pr1* KO mice using the Colony Stimulating Factor 1 Receptor (CSF1R) inhibitor PLX5622 prior to NMDA treatment and collected retinas at 2 DPI (supplemental Fig.1). Cell death was not detected in saline-injected retinas of PLX-treated animals, but we observed significant increases in numbers of TUNEL-positive cells in the INL and GCL in NMDA-treated retinas at 2 days after treatment (supplemental Fig.1). In PLX-treated animals, numbers of TUNEL+ cells were significantly reduced in both the GCL and the INL with MG-specific cKO of S1pr1 (supplemental Fig.1). Collectively, these findings indicate that S1pr1-signaling regulates cell survival following NMDA damage, and this process operates independently of signals from immune cells.

## Discussion

### Patterns of expression of S1pr1 and Sphk1 in the retinas of different vertebrates

We recently described conserved patterns of expression of S1P-related genes in the retinas of chicks, zebrafish, and humans (Taylor et al., 2024b). In the current study, we find that patterns of expression of S1P-related genes in the mouse retina are distinctly different from those seen in other vertebrates. For example, in chick, human, and zebrafish retinas, *S1pr1* is highly expressed by resting MG and is downregulated in activated MG in damaged retinas (Taylor et al., 2024b). However, in mouse and human retinas *Sphk1* is upregulated by activated glia, whereas in the chick retina *SPHK1* is downregulated in activated MG and MGPCs (Taylor et al., 2024b). It is possible that S1pr1-signaling in MG in resting and damaged retinas differ between mammals. It is possible that activation of S1pr1 “primes” or “kick-starts” pro-inflammatory responses in MG and these responses are needed to begin the process of dedifferentiation and reprogramming in chick, and perhaps zebrafish. By contrast, in mouse MG this pro-inflammatory process, which current evidence suggests involves the rapid activation of S1pr1 and S1pr3, leads to reactivity or drives MG to return to a resting phenotype (Hoang et al., 2020)

This study is the first to target Sphk1 or S1pr1 specifically in MG. Another group carried out a neuron-specific deletion of S1pr1 in an experimental model of chronic glaucoma (Basavarajappa et al., 2023a). Contrary to our report, this study observed significant thinning of the GCL and lower GCL response amplitudes in AAV-GFP-Cre:S1pr1^-/-^ mice relative to control retinas. While this group did not provide evidence that S1pr1 is expressed in retinal neurons, some studies have reported S1P receptor antibody labeling in neural cells and RPE (Ahmed et al., 2024; Joly et al., 2017; Nakamura et al., 2021; Porter et al., 2018). In the current study, we did not observe plausible patterns of labeling for Sphk1 or S1pr1 antibodies in the mouse retina. However, we do find that *S1pr1* mRNA appears predominantly in the MG in normal and NMDA-damaged retinas. Indeed, bulk RNA-seq libraries of sorted healthy brain cells indicate that *S1pr1* is highly expressed by astrocytes and, at lower levels, by endothelial cells in mouse and human (Zhang et al., 2014). A recent report using a novel S1pr1 GFP reporter mouse treated with cuprizone did not observe expression in mature neurons or astrocytes but rather in oligodendrocytes, oligodendrocyte progenitors, a subset of neural stem cells, and myeloid cells (Hashemi et al., 2023). It is possible that neural S1pr1 expression can be induced by some models of retinal damage and not others, similar to the dynamic expression of S1P receptors the CNS in response to different models of multiple sclerosis (Hashemi et al., 2023; Liu et al., 2016; Van Doorn et al., 2010).

### S1P-signaling and neuroprotection

In the current study we provide evidence that the activity of S1pr1 and Sphk1 in MG regulates neural survival and this process is not dependent on the presence of microglia. However, the mechanisms underlying MG-mediated neuronal survival are not well understood. MG span the entire width of the retina and serve many homeostatic functions, including neurotransmitter metabolism, potassium ion buffering, and inner blood brain barrier maintenance (Izumi et al., 1999; Karwoski et al., 1989; Kugler et al., 2021); it is likely that these functions change following retinal damage and following activation of different cell signaling pathways. In response to retinal injury, MG secrete neurotrophic factors to directly promote neural survival (reviewed by Eastlake et al., 2020), and factors to regulate the activity of microglia (Campbell et al., 2022; Hu et al., 2021; Xu et al., 2024). MG are known to produce different pro-survival factors that influence the survival of neurons in different models of retinal disease (Fu et al., 2015; Gao et al., 2025). It seems likely that S1pr1 activation in MG leads to changes in the production of secreted factors that impact neuronal survival.

We found that a small molecule agonist to S1pr1 activated MAPK-signaling in MG, whereas exogenous S1P did not. It is possible that the exogenous S1P was, in part, degraded by Sgpl1 and thereby did not reach sufficient nor sustained levels within the retina to activate S1pr1 and MAPK-signaling. Alternatively, different receptor-binding affinities of S1P (0.1-1uM) and SEW2871 (13 nM) may underlie the differential activation MAPK in MG. However, exogenous S1P did activate NFκB-signaling in MG, suggesting that levels of S1P were sufficient to activate this pathway. It has been reported that S1P acts intracellularly by direct interactions with TNFR-associated factor 2 (TRAF2), to recruit IκB kinase and subsequently activate NFκB (Alvarez et al., 2010; Park et al., 2015). However, high concentrations of exogenous S1P are required to significantly increase intracellular levels; consequently, relatively low concentrations of exogenous S1P stimulate MAPK but *not* NFκB (Alvarez et al., 2010; Van Brocklyn et al., 1998). It is also possible that exogenous S1P was sufficient to transiently activate MAPK and NFkB pathways, but we detected only the NFkB response because of the perdurance of the GFP reporter and rapid inactivation of pERK1/2 via DUSP1 and DUSP6 phosphatases (Kelly et al., 2024). Thus, a combination of factors, such as receptor binding affinity, intracellular activity, GFP reporter stability and sustained receptor activation, may underly the observed difference in the ability of exogenous S1P and SEW2871 to activate MAPK.

In mice, there is substantial evidence that elevated retinal S1P contributes to the degeneration of multiple retinal neuron types (Shiwani et al., 2021). In the current study, we find neuroprotective effects of inhibiting Sphk1 and or cKO this enzyme in MG in damaged retinas. However, neuroprotective effects of inhibiting S1pr1 were not observed, whereas cKO of this gene in MG in NMDA damaged retinas. These differences in numbers of dying cells might be attributed to the different timepoints at which retinas were harvested (24 HPI vs. 48 HPI), or due to the limited inhibition achieved by the drug. It is also possible that changes in cell death in drug-treated retinas could be mediated, in part, by actions on endothelial cells which express Sphk1 and S1P receptors. Finally, it is possible that a combination of factors was responsible for these discrepancies.

### S1P-signaling and the accumulation of immune cells in damaged retinas

The activation of microglia and MG in the retina is intimately linked. For example, we recently reported that hundreds of genes are up- or downregulated in MG in normal and damaged retinas when microglia are ablated in the chick retina (El-Hodiri et al., 2023). Reactive microglia and recruitment of monocytes into damaged retinas suppress the reprogramming of Ascl1-OE MG (Blasdel et al., 2024; Todd et al., 2020). Activated microglia/macrophage provide pro-inflammatory cytokines that activate NFκB-signaling in MG and, thereby, suppress Ascl1-mediated reprogramming of MG into neuron-like cells (Palazzo et al., 2022). cKO of *Ikkb* in MG, which is expected to block NFκB-signaling, or application of NFκB-inhibitor resulted in diminished accumulation of microglia/macrophage in damaged retinas, and this might result because of downregulation of *Ccl2* (Palazzo et al., 2022). We report here that cKO of *S1pr1*, but not *Sphk1*, from MG results in the accumulation of significantly more immune cells in damaged retinas, and this accumulation results, in part, from enhanced recruitment of peripheral monocytes.

We have previously reported that NFκB-signaling in MG promotes the accumulation of reactive immune cells, decreases neuronal survival, and suppresses the reprogramming of Ascl1-overexpressing MG (Palazzo et al., 2022). Many secreted factors stimulate NFκB, including proinflammatory cytokines, TNF, CNTF, and osteopontin (Ji et al., 2022; Palazzo et al., 2023). In the current study, we report that S1pr1 activation stimulates NFκB-signaling in MG and S1pr1 inhibition suppresses NFκB-signaling in the damaged retina. We propose that the S1pr1:NFκB-signaling in MG is a significant driver of ganglion cell death in NMDA-damaged retinas. This signaling relationship appears to be bidirectional, wherein MG-specific *Ikkb* knockout increases *S1pr1* expression and decreases *S1pr3* and *Sphk1* expression.

It seems inconsistent that both *Ikkb* cKO (blocked NFκB-signaling) and *S1pr1* cKO (diminished NFκB-signaling) promote neuron survival, while *Ikkb c*KO *suppresses,* whereas *S1pr1* cKO *promotes* the accumulation of immune cells in damaged retinas. Thus, NFκB-signaling in MG significantly contributes to the MG:microglia coordination, and S1pr1 may impact this coordination via pathways other than NFκB.

### S1pr1 inflammatory signaling and neurogenesis

Cytokine-mediate pro-inflammatory signaling is required to stimulate MG to become activated and proliferate as MGPCs in the retinas of chicks and zebrafish (Fischer et al., 2014; Silva et al., 2020). However, sustained microglial reactivity, cytokine signaling and NFκB activation suppress MG reprogramming (Palazzo et al., 2020; White et al., 2017a). In the current study, we provide evidence that S1P-signaling suppresses Ascl1-mediated reprogramming of MG into progenitor cells that produce bipolar-like neurons. This effect is consistent with S1pr1/3-signaling, in part, through the NFκB pathway, which is known to suppress Ascl1-mediated reprogramming of MG (Palazzo et al., 2022). However, it is likely that cell signaling pathways in addition to NFκB are impacted by activation of S1pr1/3 in MG in damaged retinas. For example, in MG in the chick retina Jak/Stat, mTOR, and Smad1/5/9 second messenger pathways are downstream of S1P-receptor activation (Taylor et al., 2024b). These cell signaling pathways are known to be activated in MG following NMDA-treatment and promote the formation of proliferating MGPCs (Todd et al., 2016; Todd et al., 2017; Zelinka et al., 2016). In the chick, S1PR1 inhibitors and NFκB inhibitors stimulate the formation of proliferating MGPCs following neuronal damage (Palazzo et al., 2020; Taylor et al., 2024b). In the absence of pro-inflammatory signals from microglia, S1PR1 *inhibitor* promotes damage-dependent MGPC formation (Taylor et al., 2025), whereas NFκB *activator* increases MGPC formation (Palazzo et al., 2020). These findings suggest that the NFκB pathway must be transiently activated to initiate reactivity and de-differentiation but thereafter must be suppressed to permit upregulation of progenitor-genes and proliferation. It is probable that signaling through S1pr1/3 potentiates or sustains NFκB-signaling to suppress reprogramming of MG into neuron-like cells.

## Conclusions

Our findings indicate that *S1pr1* and *Sphk1* are rapidly and transiently upregulated by MG following acute retinal injury in the mouse. Treatments that activate/inhibit S1P-receptors result in increased/decreased NFκB-signaling in MG in normal and damaged retinas. The expression of *S1pr1* in MG is upregulated by NFκB-signaling and downregulated by forced expression of Ascl1. Selective deletion of *S1pr1* and/or *Sphk1* from MG has no apparent effects on undamaged retinas, whereas the deletion of these genes from MG in damaged retinas increases the accumulation of immune cells, decreases cell death and increases neuronal survival of inner retinal neurons. Consistent with these findings, pharmacological treatments that increase S1P-signaling increased cell death, whereas treatments that decreased S1P-signaling decreased cell death in damaged retinas. We conclude that the inflammation mediated by autocrine signaling via S1pr1 in MG in damaged retinas suppresses the neuroprotective functions of MG and suppress Ascl1-mediated reprogramming of MG into neuron-like cells.

## Data availability

CellRanger output files for Gene-Cell matrices for scRNA-seq data for libraries from saline and NMDA-treated retinas are available through Sharepoint links for embryonic chick retina databases: https://osumc.sharepoint.com/:f:/s/Links/Eoto-Qg2uuxDn1bHWMM6gdkBTft4S_YSBjQJResxY-qehA?e=mdaPlg and post-hatch normal and treated chick retina databases: https://osumc.sharepoint.com/:f:/s/Links/Eoto-Qg2uuxDn1bHWMM6gdkBTft4S_YSBjQJResxY-qehA?e=mdaPlg. scRNA-seq datasets are deposited in GEO (GSE135406, GSE242796) and Gene-Cell matrices for scRNA-seq data for libraries chick retinas treated with saline or NMDA retinas are available through NCBI (GSM7770646, GSM7770647, GSM7770648, GSM7770649).

Gene-cell matrices for scRNA-seq data are deposited at GitHub: scRNA-seq data from NMDA-damaged mouse retinas (Hoang et al., 2020) can be queried here: https://proteinpaint.stjude.org/F/2019.retina.scRNA.html

ScRNA-seq data for mouse retina at different stage of post-natal development (Li et al., 2024, iScience) https://cellxgene.cziscience.com/collections/a0c84e3f-a5ca-4481-b3a5-ccfda0a81ecc

## Author Contributions

OT designed and executed experiments, gathered data, constructed figures and wrote the manuscript. LK executed experiments and gathered data. AJF designed experiments, analyzed data, constructed figures and wrote the manuscript. HE wrote the manuscript.

## Acknowledgements

We wish to thank Timothy Hla for providing the *Sphk1*^f/f^ and *S1pr1* ^f/f^ mice. We also thank Dr. Ed Levine for providing the Rlbp1-CreERT mice and Dr. Dennis Guttridge for providing the NFκB-eGFP reporter mice.

---

**Supplemental Figure 1: cKO of S1PR1 is neuroprotective in the absence of retinal immune cells**

Rlbp1-CreERT:S1pr1fl/fl mice were fed PLX5622 chow ad libitum for two weeks, then IP injected with tamoxifen or corn oil once daily for 4 days. Eyes were intravitreally injected with NMDA or saline control, and retinas were collected 48HPI. Sections of the retina were labeled for fragmented DNA (TUNEL; green) and DAPI (blue; a). Arrows indicate TUNEL+ puncta. Calibration bar in panel a represents 50 µm. Histograms illustrate mean (bar ± SD) numbers of TUNEL+ cells in the INL and ONL (b) and GCL (c). Each dot represents one biological replicate. Significance of difference (p-values) was determined by using ANOVA with Sidak correction. Abbreviations: ONL, outer nuclear layer; INL, inner nuclear layer; IPL, inner plexiform layer; GCL, ganglion cell layer; NMDA, N-methyl-D-aspartate; ns, not significant; TUNEL, terminal deoxynucleotidyl transferase dUTP nick end labeling.

