## Supplemental Table 1 for "Sphingosine-1-phosphate signaling through Müller glia regulates neuroprotection and the accumulation of immune cells in the rodent retina"

**Figure 2j:** Statistics for expression of S1pr1, S1pr3, Sphk1, Vim, Tgfb2 and Klf6 in Muller glia – retina atlas.

ENSMUSG00000045092 – S1pr1

ENSMUSG00000067586 – S1pr3

ENSMUSG00000061878 – Sphk1

ENSMUSG00000026728 – Vim

ENSMUSG00000039239 – Tgfb2

ENSMUSG00000000078 – Klf6

```
FindMarkers(MG_ret_atlas, ident.1 = c("2-week-old stage"), ident.2 = c("5-wee
k-old stage", "7-week-old stage", "8-week-old stage",  "9-week-old stage", "1
1-week-old stage", "12-week-old stage", "4-month-old stage", "25-week-old sta
ge"), group.by = "development_stage", features = c("ENSMUSG00000045092", "ENS
MUSG00000067586", "ENSMUSG00000061878", "ENSMUSG00000026728", "ENSMUSG0000000
0078"), min.pct = 0.001, logfc.threshold = 0.001, test.use = "wilcox")
                          p_val avg_log2FC pct.1 pct.2    p_val_adj
ENSMUSG00000026728 2.739862e-48  0.9454207 0.950 0.848 8.776874e-44
ENSMUSG00000061878 1.177297e-08  0.9471557 0.306 0.177 3.771354e-04
ENSMUSG00000045092 1.531962e-05 -0.7244541 0.575 0.612 4.907487e-01
ENSMUSG00000067586 7.972875e-04 -0.7865178 0.442 0.477 1.000000e+00
ENSMUSG00000000078 9.503804e-02 -1.1028531 0.236 0.251 1.000000e+00
```

```
FindMarkers(MG_ret_atlas, ident.1 = c("4-month-old stage", "25-week-old stage
"), ident.2 = c("2-week-old stage", "5-week-old stage", "7-week-old stage", "
9-week-old stage", "11-week-old stage", "12-week-old stage"), group.by = "dev
elopment_stage", features = c("ENSMUSG00000045092", "ENSMUSG00000067586", "EN
SMUSG00000061878", "ENSMUSG00000026728", "ENSMUSG00000000078"), min.pct = 0.0
01, logfc.threshold = 0.001, test.use = "wilcox")
                          p_val avg_log2FC pct.1 pct.2    p_val_adj
ENSMUSG00000026728 1.911385e-35 -0.6495088 0.764 0.894 6.122932e-31
ENSMUSG00000045092 5.105467e-13  0.8520706 0.654 0.522 1.635485e-08
ENSMUSG00000067586 4.860333e-07  0.6580542 0.454 0.340 1.556959e-02
ENSMUSG00000061878 5.399536e-05 -1.2261121 0.106 0.177 1.000000e+00
ENSMUSG00000000078 3.609386e-02 -0.4152630 0.115 0.153 1.000000e+00
```


```
FindMarkers(MG_ret_atlas, ident.1 = c("4-month-old stage", "25-week-old stage
"), ident.2 = c("2-week-old stage", "5-week-old stage", "7-week-old stage", "
9-week-old stage", "11-week-old stage", "12-week-old stage"), group.by = "dev
elopment_stage", features = c("ENSMUSG00000045092", "ENSMUSG00000067586", "EN
SMUSG00000061878", "ENSMUSG00000026728", "ENSMUSG00000000078"), min.pct = 0.0
01, logfc.threshold = 0.001, test.use = "wilcox")
                          p_val avg_log2FC pct.1 pct.2    p_val_adj
ENSMUSG00000026728 1.911385e-35 -0.6495088 0.764 0.894 6.122932e-31
ENSMUSG00000045092 5.105467e-13  0.8520706 0.654 0.522 1.635485e-08
ENSMUSG00000067586 4.860333e-07  0.6580542 0.454 0.340 1.556959e-02
ENSMUSG00000061878 5.399536e-05 -1.2261121 0.106 0.177 1.000000e+00
ENSMUSG00000000078 3.609386e-02 -0.4152630 0.115 0.153 1.000000e+00
```

```
FindMarkers(MG_ret_atlas, ident.1 = c("8-week-old stage"), ident.2 = c("2-wee
k-old stage", "5-week-old stage", "7-week-old stage", "9-week-old stage", "11
-week-old stage", "12-week-old stage"), group.by = "development_stage", featu
res = c("ENSMUSG00000045092", "ENSMUSG00000067586", "ENSMUSG00000061878", "EN
SMUSG00000026728", "ENSMUSG00000000078"), min.pct = 0.001, logfc.threshold =
0.001, test.use = "wilcox")
                          p_val avg_log2FC pct.1 pct.2    p_val_adj
ENSMUSG00000026728 9.758767e-67 -0.5829743 0.852 0.894 3.126123e-62
ENSMUSG00000067586 6.986148e-31  0.8586471 0.508 0.340 2.237943e-26
ENSMUSG00000000078 5.212162e-26  1.7467857 0.287 0.153 1.669664e-21
ENSMUSG00000045092 7.010688e-10  0.3281378 0.625 0.522 2.245804e-05
ENSMUSG00000061878 4.504073e-01 -0.1025473 0.191 0.177 1.000000e+00
```

**Figure 3i**: Statistics mouse Muller glia – S1P related genes in Muller glia

```
FindMarkers(mouse_MG, ident.1 = "ctrl MG", ident.2 = "3hr MG", features = c("
S1pr1", "S1pr2", "S1pr3", "Sphk1", "Sgpl1"), min.pct = 0.001, logfc.threshold
= 0.001, test.use = "poisson")
  |++++++++++++++++++++++++++++++++++++++++++++++++++| 100% elapsed=00s
             p_val  avg_log2FC pct.1 pct.2    p_val_adj
Sphk1 0.000000e+00 -1.61452559 0.124 0.727 0.000000e+00
S1pr1 5.620102e-50 -0.33240797 0.254 0.356 1.014147e-45
Sgpl1 1.056373e-11 -0.08021554 0.039 0.092 1.906225e-07
S1pr3 3.523249e-10 -0.10989705 0.131 0.180 6.357703e-06
S1pr2 3.126410e-08 -0.04930048 0.017 0.052 5.641607e-04

FindMarkers(mouse_MG, ident.1 = "ctrl MG", ident.2 = "6hr MG", features = c("
S1pr1", "S1pr2", "S1pr3", "Sphk1", "Sgpl1"), min.pct = 0.001, logfc.threshold
= 0.001, test.use = "poisson")
  |++++++++++++++++++++++++++++++++++++++++++++++++++| 100% elapsed=00s
              p_val  avg_log2FC pct.1 pct.2     p_val_adj
Sphk1  0.000000e+00 -1.56261852 0.124 0.744  0.000000e+00
S1pr1 7.438025e-291 -1.03865566 0.254 0.631 1.342192e-286
Sgpl1  4.205724e-18 -0.12441949 0.039 0.119  7.589229e-14
S1pr3  1.243471e-04 -0.07420322 0.131 0.155  1.000000e+00
S1pr2  1.707192e-04 -0.03339105 0.017 0.041  1.000000e+00

FindMarkers(mouse_MG, ident.1 = "ctrl MG", ident.2 = "12+24hr MG", features =
c("S1pr1", "S1pr2", "S1pr3", "Sphk1", "Sgpl1"), min.pct = 0.001, logfc.thresh
old = 0.001, test.use = "poisson")
  |++++++++++++++++++++++++++++++++++++++++++++++++++| 100% elapsed=00s
             p_val  avg_log2FC pct.1 pct.2    p_val_adj
S1pr1 7.212750e-34 -0.34226544 0.254 0.269 1.301541e-29
Sphk1 1.573449e-14 -0.17414569 0.124 0.189 2.839289e-10
Sgpl1 2.104924e-04 -0.05054016 0.039 0.068 1.000000e+00
S1pr2 1.002914e-02 -0.02478273 0.017 0.033 1.000000e+00
S1pr3 6.281689e-02  0.04028247 0.131 0.088 1.000000e+00

FindMarkers(mouse_MG, ident.1 = "ctrl MG", ident.2 = "36+48+72hr MG", feature
s = c("S1pr1", "S1pr2", "S1pr3", "Sphk1", "Sgpl1"), min.pct = 0.001, logfc.th
reshold = 0.001, test.use = "poisson")
  |++++++++++++++++++++++++++++++++++++++++++++++++++| 100% elapsed=00s
             p_val  avg_log2FC pct.1 pct.2    p_val_adj
S1pr3 4.794872e-83 -0.56672067 0.131 0.417 8.652346e-79
S1pr1 1.603664e-16 -0.25855050 0.254 0.392 2.893812e-12
Sgpl1 1.710830e-11 -0.11432589 0.039 0.117 3.087193e-07
S1pr2 4.474202e-07 -0.06261198 0.017 0.060 8.073697e-03
Sphk1 3.489986e-01  0.02228651 0.124 0.115 1.000000e+00
```

```
FindMarkers(mouse_MG, ident.1 = "12+24hr MG", ident.2 = "3hr MG", features =
c("S1pr1", "S1pr2", "S1pr3", "Sphk1", "Sgpl1"), min.pct = 0.001, logfc.thresh
old = 0.001, test.use = "poisson")
  |++++++++++++++++++++++++++++++++++++++++++++++++++| 100% elapsed=00s
              p_val   avg_log2FC pct.1 pct.2     p_val_adj
Sphk1 2.624011e-195 -1.44037989 0.189 0.727 4.735028e-191
S1pr3  4.270799e-09 -0.15017951 0.088 0.180  7.706657e-05
S1pr2  7.265473e-02 -0.02451775 0.033 0.052  1.000000e+00
Sgpl1  9.877888e-02 -0.02967539 0.068 0.092  1.000000e+00
S1pr1  7.526442e-01  0.00985747 0.269 0.356  1.000000e+00
> FindMarkers(mouse_MG, ident.1 = "12+24hr MG", ident.2 = "6hr MG", features
= c("S1pr1", "S1pr2", "S1pr3", "Sphk1", "Sgpl1"), min.pct = 0.001, logfc.thre
shold = 0.001, test.use = "poisson")
  |++++++++++++++++++++++++++++++++++++++++++++++++++| 100% elapsed=00s
              p_val    avg_log2FC pct.1 pct.2     p_val_adj
Sphk1 8.847884e-181 -1.388472822 0.189 0.744 1.596601e-176
S1pr1  3.633764e-73 -0.696390220 0.269 0.631  6.557127e-69
S1pr3  8.878377e-06 -0.114485685 0.088 0.155  1.602103e-01
Sgpl1  5.320239e-04 -0.073879331 0.068 0.119  1.000000e+00
S1pr2  5.207173e-01 -0.008608314 0.033 0.041  1.000000e+00

FindMarkers(mouse_MG, ident.1 = "36+48+72hr MG", ident.2 = "12+24hr MG", feat
ures = c("S1pr1", "S1pr2", "S1pr3", "Sphk1", "Sgpl1"), min.pct = 0.001, logfc
.threshold = 0.001, test.use = "poisson")
  |++++++++++++++++++++++++++++++++++++++++++++++++++| 100% elapsed=00s
             p_val  avg_log2FC pct.1 pct.2    p_val_adj
S1pr3 1.277570e-47  0.60700313 0.417 0.088 2.305374e-43
Sphk1 8.578326e-09 -0.19643220 0.115 0.189 1.547959e-04
Sgpl1 8.803970e-03  0.06378573 0.117 0.068 1.000000e+00
S1pr2 3.510023e-02  0.03782924 0.060 0.033 1.000000e+00
S1pr1 4.560113e-02 -0.08371494 0.392 0.269 1.000000e+00

FindMarkers(mouse_MG, ident.1 = "36+48+72hr MG", ident.2 = "6hr MG", features
= c("S1pr1", "S1pr2", "S1pr3", "Sphk1", "Sgpl1"), min.pct = 0.001, logfc.thre
shold = 0.001, test.use = "poisson")
  |++++++++++++++++++++++++++++++++++++++++++++++++++| 100% elapsed=00s
              p_val   avg_log2FC pct.1 pct.2     p_val_adj
Sphk1 7.157446e-113 -1.58490502 0.115 0.744 1.291561e-108
S1pr1  5.990276e-65 -0.78010516 0.392 0.631  1.080945e-60
S1pr3  4.621340e-46  0.49251745 0.417 0.155  8.339208e-42
S1pr2  7.658780e-02  0.02922093 0.060 0.041  1.000000e+00
Sgpl1  6.868297e-01 -0.01009360 0.117 0.119  1.000000e+00

FindMarkers(mouse_MG, ident.1 = "36+48+72hr MG", ident.2 = "3hr MG", features
= c("S1pr1", "S1pr2", "S1pr3", "Sphk1", "Sgpl1"), min.pct = 0.001, logfc.thre
shold = 0.001, test.use = "poisson")
  |++++++++++++++++++++++++++++++++++++++++++++++++++| 100% elapsed=00s
              p_val   avg_log2FC pct.1 pct.2     p_val_adj
Sphk1 8.422922e-118 -1.63681209 0.115 0.727 1.519916e-113
S1pr3  1.193114e-50  0.45682362 0.417 0.180  2.152974e-46
S1pr1  4.203181e-02 -0.07385747 0.392 0.356  1.000000e+00
Sgpl1  1.091430e-01  0.03411035 0.117 0.092  1.000000e+00
S1pr2  4.155368e-01  0.01331149 0.060 0.052  1.000000e+00
```


```
FindMarkers(mouse_MG, ident.1 = "12+24hr MG", ident.2 = "3hr MG", features =
c("S1pr1", "S1pr2", "S1pr3", "Sphk1", "Sgpl1"), min.pct = 0.001, logfc.thresh
old = 0.001, test.use = "poisson")
  |++++++++++++++++++++++++++++++++++++++++++++++++++| 100% elapsed=00s
              p_val   avg_log2FC pct.1 pct.2     p_val_adj
Sphk1 2.624011e-195 -1.44037989 0.189 0.727 4.735028e-191
S1pr3  4.270799e-09 -0.15017951 0.088 0.180  7.706657e-05
S1pr2  7.265473e-02 -0.02451775 0.033 0.052  1.000000e+00
Sgpl1  9.877888e-02 -0.02967539 0.068 0.092  1.000000e+00
S1pr1  7.526442e-01  0.00985747 0.269 0.356  1.000000e+00
```

```
FindMarkers(mouse_MG, ident.1 = "12+24hr MG", ident.2 = "6hr MG", features =
c("S1pr1", "S1pr2", "S1pr3", "Sphk1", "Sgpl1"), min.pct = 0.001, logfc.thresh
old = 0.001, test.use = "poisson")
  |++++++++++++++++++++++++++++++++++++++++++++++++++| 100% elapsed=00s
              p_val   avg_log2FC pct.1 pct.2     p_val_adj
Sphk1 8.847884e-181 -1.388472822 0.189 0.744 1.596601e-176
S1pr1  3.633764e-73 -0.696390220 0.269 0.631  6.557127e-69
S1pr3  8.878377e-06 -0.114485685 0.088 0.155  1.602103e-01
Sgpl1  5.320239e-04 -0.073879331 0.068 0.119  1.000000e+00
S1pr2  5.207173e-01 -0.008608314 0.033 0.041  1.000000e+00

FindMarkers(mouse_MG, ident.1 = "3hr MG", ident.2 = "6hr MG", features = c("S
1pr1", "S1pr2", "S1pr3", "Sphk1", "Sgpl1"), min.pct = 0.001, logfc.threshold
= 0.001, test.use = "poisson")
  |++++++++++++++++++++++++++++++++++++++++++++++++++| 100% elapsed=00s
              p_val  avg_log2FC pct.1 pct.2     p_val_adj
S1pr1 2.659847e-153 -0.70624769 0.356 0.631 4.799694e-149
Sgpl1  6.423643e-03 -0.04420395 0.092 0.119  1.000000e+00
Sphk1  3.507069e-02  0.05190707 0.727 0.744  1.000000e+00
S1pr3  9.462036e-02  0.03569383 0.180 0.155  1.000000e+00
S1pr2  1.665002e-01  0.01590944 0.052 0.041  1.000000e+00
```
