## Supplementary figures and images for "Sphingosine-1-phosphate signaling through Müller glia regulates neuroprotection and the accumulation of immune cells in the rodent retina"

### supplemental Figure 1

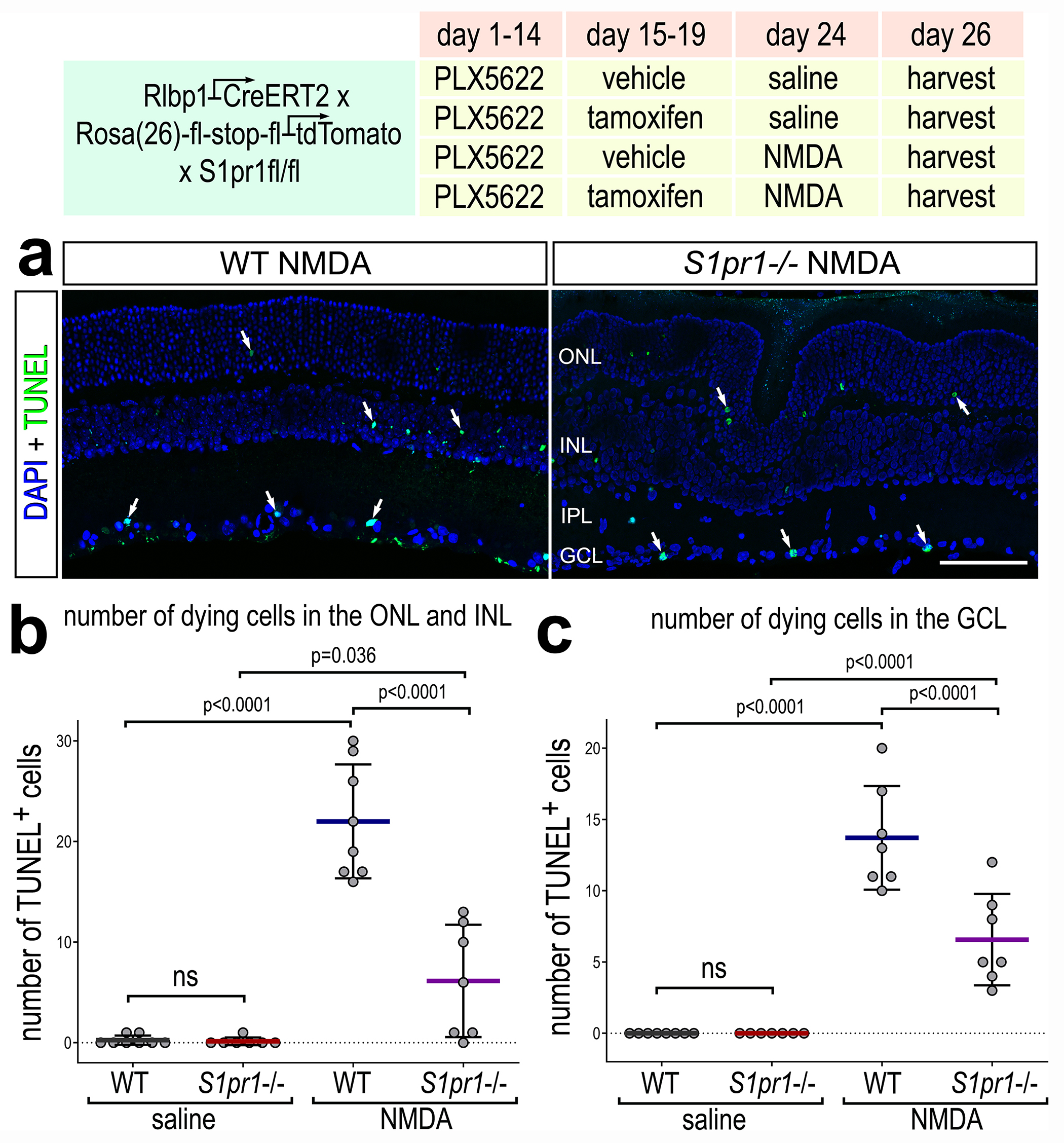
